# In silico designed small molecule targets misfolded TDP-43 to rescue function and ALS pathology

**DOI:** 10.64898/2026.11.14.766688

**Authors:** Marcela Kokes, Devi Supriya Korumilli, Vanessa Seawright, Shruti Arya, Annika van Hummel, Miheer Sabale, Md. Anzarul Haque, Naomi Moreno, Agenor Limon, Pritam Ganguly, Eric Shao, Yazi D. Ke, Rakez Kayed, Joan-Emma Shea, Ambuj Singh, Lewis Whitehead, Catherine Planey, Vidhu Mathur

## Abstract

The DNA/RNA binding protein TARDBP or TDP-43 is a key driver of pathology in Amyotrophic Lateral Sclerosis (ALS), a motor neuron disease with no known cure. In >97% of sporadic ALS patients, nuclear TDP-43 in motor neurons mislocalizes and aggregates in the cellular cytoplasm (gain-of-function). A parallel reduction in nuclear TDP-43 in ALS patients leads to a loss-of-function in mRNA binding and splicing regulation. Using our machine learning platform, we discovered ACE-339, a novel, orally bioavailable small molecule that disrupts insoluble TDP-43 aggregates. Using *in vitro*, *in vivo* and *in silico* models, we show that ACE-339 directly targets aggregated forms of TDP-43. ACE-339 interacts directly with TDP-43 hexamers, disaggregates recombinant TDP-43 and rescues both gain- and loss-of-function phenotypes of TDP-43 in cells. ACE-339 functionally rescues TDP-43 dependent mis-splicing and synaptic activity. Lastly, ACE-339 rescues disease associated phenotypes in two independent mouse models of TDP-43 pathology. In addition, we now have a catalog of TDP-43 disaggregating compounds that are being developed further to be tested in clinical trials in patients with ALS and other TDP-43 proteinopathies.

## Introduction

TAR DNA-binding protein or TDP-43 is a ubiquitously expressed hnRNP family member that regulates critical cellular processes including RNA metabolism, mRNA splicing and transport, microRNA maturation, and the repression of cryptic exons^1,2^. TDP-43 also plays a critical role in the development of the central nervous system (CNS)^3^. Increasingly, TDP-43 dysfunction is recognized as a critical pathogenic mechanism across neurodegenerative diseases.

Many neuromuscular and neurodegenerative diseases, collectively called TDP-43 proteinopathies, present with TDP-43 loss-of-function and gain of function phenotypes^2,4^. Almost all sporadic Amyotrophic Lateral Sclerosis (ALS) and Frontotemporal Lobar Degeneration (FTLD-TDP) patients display TDP-43 nuclear loss and cytoplasmic aggregates in the CNS^5,6^. Additionally, individuals with TDP-43 mutations that enhance these cellular phenotypes develop ALS/FTLD with an accelerated disease pathology^7–10^. Therefore, targeting disease associated TDP-43 phenotypes could potentially reverse neuronal toxicity and delay the progression of diseases.

An important pathological feature of ALS and FTLD-TDP is the cytoplasmic accumulation of insoluble, hyperphosphorylated and ubiquitinated TDP-43 inclusions^11,12^. TDP-43 shuttles between the cytoplasm and nucleus but is predominantly nuclear under physiological conditions, where it enacts its critical functions including RNA splicing regulation. In diseases, TDP-43 exhibits a dual pathology: “loss-of-function” due to the loss of nuclear, functional TDP-43 and “gain-of-function” due to the accumulation of misfolded, cytoplasmic TDP-43 inclusions^13–15^. Importantly, cytoplasmic aggregation was shown to be sufficient to drive TDP-43 nuclear clearance^16^, stripping the cell of critical splicing regulation. The loss of nuclear TDP-43 leads to the aberrant inclusion of cryptic exons in essential neuronal transcripts, such as STMN2 and UNC13A – a molecular hallmark of ALS and FTD that ultimately drives neuronal dysfunction and cell death.

TDP-43 functions via its structured N-terminal domain which contains the nuclear localization signal and two RNA recognition motifs and forms functional homodimers^17^. The C-terminal low complexity domain (LCD) is disordered and contains prion-like gln/asn-rich and gly-rich regions^18,19^. Through the LCD, TDP-43 assembles into amyloid-like aggregates in the cytoplasm, rendering it non-functional and contributing to disease pathogenesis. Multiple mechanisms including post-translational modifications, nucleocytoplasmic transport dysfunction, liquid-liquid phase separation and autophagy have been suggested to trigger TDP-43 oligomerization and further aggregation in the cytoplasm^20^.

Here we present ACE-339, a small molecule identified using the RESOLUTE AI-driven computational chemistry platform which is designed to discover drug-like chemical matter, elucidate protein-protein interactions, and identify cryptic pockets. We show that ACE-339 directly targets the misfolded, insoluble, gain-of-function phenotype of TDP-43 (henceforth called “aggregates”): ACE-339 disrupts recombinant TDP-43 aggregates, cytoplasmic TDP-43 puncta and insoluble TDP-43 in the brains of a mouse model. Molecular dynamics simulations further support the direct activity of ACE-339 on TDP-43 aggregates upon engagement with a potential cryptic pocket. We further show that this activity of ACE-339 leads to a rescue of the loss-of-function phenotype of TDP-43: ACE-339 restores nuclear TDP-43 and rescues TDP-43-dependent mis-splicing and synaptic dysfunction. ACE-339 also improves survival, lowers neuroinflammation and Nf-L, and improves behavioral phenotypes in TDP-43 pathology-dependent mouse models. The data presented here demonstrate the capabilities of the RESOLUTE platform by successfully designing small molecules (like ACE-339) that disrupt TDP-43 aggregation to rescue gain-of-function and loss-of-function phenotypes in TDP-43 proteinopathies.

## Results

### Identification of ACE-339 through integrated *in silico* and phenotypic screening

We identified ACE-339, a small molecule that targets aggregated TDP-43, using our RESOLUTE AI-driven computational chemistry platform. The foundational RESOLUTE algorithm incorporated a Joint Pharmacophore Space (JPS) model^21^ that was previously used to discover small molecule inhibitors of LRRK2^22^ and amyloid beta oligomer formation^23^. Given the biological complexity of TDP-43 and lack of publicly available previous chemical matter directly targeting aggregated TDP-43, the RESOLUTE platform was further developed to incorporate protein-protein interaction modeling and physical *in vitro* mechanistic assays to produce a TDP-43 aggregate structure-activity relationship model. The JPS model calculates the three-dimensional molecular characteristics of virtually generated molecules and their pharmacophoric features to virtually screen chemical compound libraries and discover novel structural starting points. Building on this validated methodology, we generated a focused library of novel compounds and developed a molecular dynamics simulation engine to evaluate the disaggregation activity of compounds on TDP-43 oligomers.

To empirically evaluate the effect of this computationally designed library on TDP-43 aggregation, we developed a dual screening approach (Figure 1A). We evaluated candidates in parallel for their ability to inhibit de novo aggregation in a cell-free Thioflavin T (ThT) fluorescence assay using the amyloidogenic TDP-43 (307-319) core peptide, and for their phenotypic activity in reducing stress-induced cytoplasmic TDP-43 puncta in a U2OS reporter cell line. Among a representative panel of candidates evaluated using the RESOLUTE platform and across these assays (Figure 1B), ACE-339 emerged as one of the most promising compounds. ACE-339 demonstrated potent activity in both biochemical disaggregation and cellular TDP-43 puncta reduction, therefore it was selected for comprehensive *in vitro* and *in vivo* characterization.

**Figure 1.**
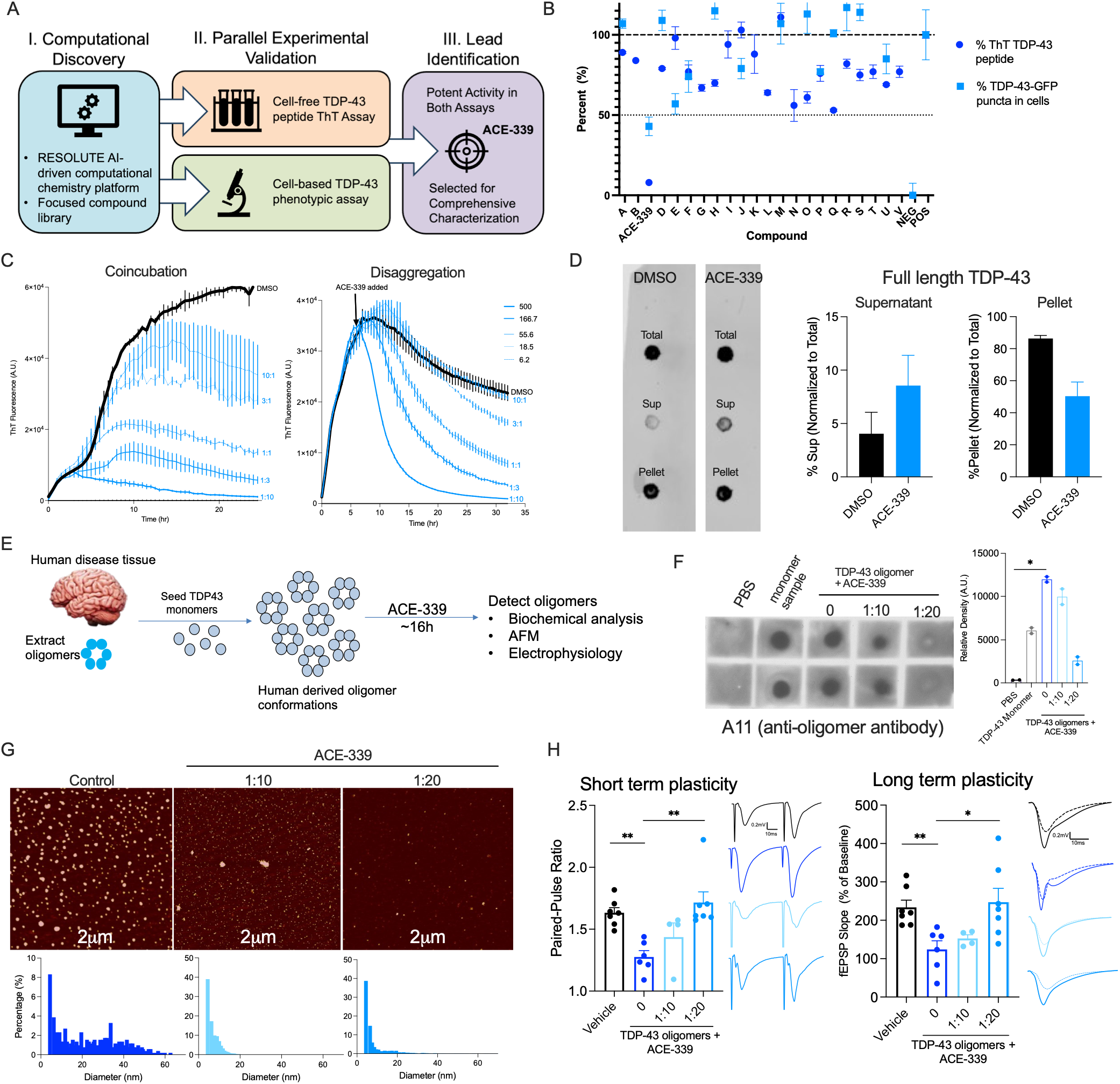
ACE-339 targets misfolded forms of TDP-43. (A) Schematic of dual screening approach. (B) Representative data from primary screen results showing candidate compound efficacy in reducing TDP-43 (307-319) peptide aggregation (dark blue) and stress-induced TDP-43-GFP cytoplasmic puncta (light blue). ACE-339 emerged as one of the lead candidates. (C) Kinetic curves of recombinant TDP-43 (307-319) core peptide showing Thioflavin T (ThT) dye binding upon coincubation (drug added at 0h) and disaggregation (drug added at 6h) with DMSO vehicle control (black) and reduced ThT fluorescence with ACE-339 at indicated concentrations (µM) (blue). Peptide monomer: drug molar ratio for each concentration are indicated. (D) ACE-339 shows increased soluble and decreased insoluble full length recombinant TDP-43 in dot blot assay where pre-formed TDP-43 aggregates were separated into supernatant and pellet by high-speed centrifugation. (E) Schematic showing experimental design for F-H. ACE-339 dissociates ALS patient brain derived oligomer seeded full length TDP-43 conformations at indicated protein to drug molar ratios. (F) ACE-339 shows reduction in A11-19 oligomer specific antibody signal seen in control, vehicle sample. Monomer sample shows A11-19 signal because it easily aggregates during experimentation. Image and quantification are representative of 3 independent runs with two technical replicates each. (G) Atomic force microscopy representative images and average size distribution of particles shows reduction in full length TDP-43 oligomers size upon ACE-339 treatment is shown. Image of 2µm x 2µm region and quantification are representative of 2 independent runs with two technical replicates each. (H) ACE-339 rescues TDP-43 oligomer induced synaptic defects in mouse hippocampal acute slice model. Quantification and representative fEPSP (dotted line=baseline, solid line=post-HFS) traces are shown (n=4-6 slices/group). Mean±SEM are shown. P<0.05*, P<0.01** One-way ANOVA followed by Dunnett’s test.

### ACE-339 specifically targets TDP-43 aggregates

TDP-43 (307-319) peptide is part of the TDP-43 amyloid core observed in filaments isolated from ALS patients and is known to aggregate independently^24,25^. In a Thioflavin T (ThT) fluorescence assay, ACE-339 dose-dependently inhibited *de novo* aggregation and disrupted pre-formed TDP-43 (307-319) aggregates (Figure 1C). Notably, the degree of inhibition at any given ACE-339 concentration was dependent on the concentration of TDP-43 (307-319) peptide in the assay (Figure S1A). Since the sensitivity of this assay necessitates using micromolar TDP-43 concentrations, the IC_50_ of ACE-339 was also in the micromolar range. We found that irrespective of the TDP-43 concentration, the stoichiometric ratio of TDP-43 (307-319) to ACE-339 required for activity was highly consistent. Accordingly, ACE-339 consistently inhibited TDP-43 aggregation >90% at 1:10, and >50% at 1:1 TDP-43 monomer to compound molar ratio at any given TDP-43 (307-319) concentration (Figure S1A) implying that molar ratios may be a more appropriate measure of potency of the compound in this assay.

ACE-339 also inhibited the aggregation of both the disease-associated low complexity domain (LCD) of TDP-43 and recombinant full length (FL) TDP-43 (Figure S1B-C) in a ThT-based assay. The shape of the ThT curves over time with ACE-339 was biphasic with ThT increasing normally at first but peaking and reducing over time, consistent with the design hypothesis that ACE-339 specifically binds to misfolded forms of TDP-43 and hence, in this assay TDP-43 aggregates first before ACE-339 exerts an effect. Importantly, ACE-339 does not absorb light at either the excitation or emission wavelengths of ThT (Figure S1D), indicating that the reduction in ThT fluorescence was not a false positive due to energy interference between ACE-339 and ThT. To further confirm the effect of ACE-339 on TDP-43 aggregates with an orthogonal dye-free method, we used high-speed centrifugation to separate soluble and aggregated FL TDP-43. Consistent with the ThT data, treatment of pre-formed FL TDP-43 aggregates with ACE-339 reduced insoluble and increased soluble TDP-43 (Figure 1D) further confirming the direct activity of ACE-339 on TDP-43 aggregates.

Since TDP-43 protein is prone to oligomerization and aggregation, we confirmed that the recombinant FL TDP-43 used in this study is initially monomeric and natively folded and becomes oligomeric and aggregates over time (Figure S1E). As expected, ACE-339 did not bind to natively folded TDP-43 (Figure S1F). We hypothesized that once ACE-339 disaggregates TDP-43, the loss of its binding site would release ACE-339 from its target protein and it would be free to continually recycle and engage with other TDP-43 aggregates. Indeed, when free and target-bound ACE-339 fractions were separated, the majority of ACE-339 was measured in the “free drug” fraction (Figure S2, top right). Moreover, the free ACE-339 in this fraction potently inhibited the aggregation of freshly added TDP-43 (307-319) (Figure S2, bottom right). We attribute the apparent higher potency in this secondary ThT assay to volume loss during the fractionation, leading to an artificially higher concentration of the compound in the free fraction. These data support a TDP-43 conformation-specific dynamic equilibrium mechanism of action for ACE-339. By specifically engaging misfolded, aggregated forms of TDP-43 and driving their dissociation, ACE-339 loses its binding site upon TDP-43 refolding, rendering it freely available to recycle and act on subsequent TDP-43 aggregates.

ACE-339 did not inhibit polymerization of unrelated, functional tubulin monomers into microtubules (Figure S1G), indicating that ACE-339 acts specifically on beta-sheet rich amyloid conformations and suggesting that it does not disrupt the native beta-sheet conformations of functional proteins. Taken together, these data demonstrate that ACE-339 specifically and directly acts on aggregated forms of TDP-43.

### ACE-339 dissociates ALS disease relevant conformations of TDP-43

To evaluate if ACE-339 can target human disease-relevant TDP-43 aggregate conformations, we purified TDP-43 oligomers from post-mortem ALS patient brains using the oligomer-specific antibody A11^26^ and used these oligomers to seed recombinant FL TDP-43 to generate amplified brain-derived TDP-43 oligomers (Figure 1E). We confirmed that these amplified oligomers maintained the distinct high molecular weight, TDP-43-positive signature of the original brain-derived material, and matured into larger amyloid fibrils, demonstrating that they represent biologically active, disease-relevant conformations (Figure S3A-C). Upon treatment with ACE-339, there was a dose-dependent reduction and complete dissolution of A11^+^ TDP-43 oligomers at 1:20 TDP-43:ACE-339 ratio (Figure 1F). Furthermore, atomic force microscopy showed that ACE-339 treatment reduced the average diameter and number of these TDP-43 oligomers (Figure 1G). The average diameter of FL TDP-43 oligomers was found to be 25.0 ±4.0nm, 15.0 ±3.0nm and 6.0 ±1.0nm in the presence of none, 1:10 and 1:20 TDP-43: ACE-339 ratios, respectively.

While amyloid-beta^27,28^, tau^29–31^ and alpha-synuclein^32,33^ oligomers have been shown to impair synaptic plasticity and transmission, this electrophysiological phenotype had not previously been demonstrated with TDP-43 oligomers. Here, we show that amplified ALS patient derived TDP-43 oligomers alone impair synaptic transmission and plasticity in mouse hippocampal slices. Importantly, ACE-339 treatment dose dependently rescued paired-pulse facilitation and long-term potentiation implying a restoration of both short- and long-term plasticity respectively in this model of TDP-43 oligomer pathology (Figure 1H and S4). Taken together, these data suggest that ACE-339 targets ALS disease relevant TDP-43 conformations, rescues TDP-43 oligomer induced synaptic defects and specifically dissociates oligomeric, pathological forms of TDP-43.

### ACE-339 rescues both gain-of-function and loss-of-function disease associated TDP-43 cellular phenotypes

To model the disease associated loss of nuclear TDP-43 and its increased deposition into cytoplasmic inclusion bodies in the brain and spinal cord neurons^1,11,12^, we used a cellular sodium arsenite stress model in a U2OS reporter cell line with inducible expression of TDP-43-GFP to quantify cytoplasmic puncta (gain-of-function) and nuclear loss (loss-of-function) of TDP-43. Previous studies showed that sodium arsenite treatment causes the formation of cytoplasmic stress granules that initially recruit TDP-43 but then TDP-43 de-mixes into solid-like punctate structures which display markers of aggregation^16,34–37^. Consistently, sodium arsenite stressed cells overexpressing TDP-43-GFP formed cytoplasmic puncta and ACE-339 reduced the number of these puncta (Figure 2A). In an orthogonal stress-free system, we found that over-expression of TDP-43-GFP alone was sufficient to form cytoplasmic puncta in a small subset of cells. ACE-339 reduced the number and frequency in the population of TDP-43-GFP puncta in this model suggesting that ACE-339 activity is independent of sodium arsenite stress (Figure S5A). Taken together, these data demonstrate that ACE-339 can inhibit the gain-of-function TDP-43 phenotype in multiple distinct cellular models of TDP-43 aggregation.

**Figure 2.**
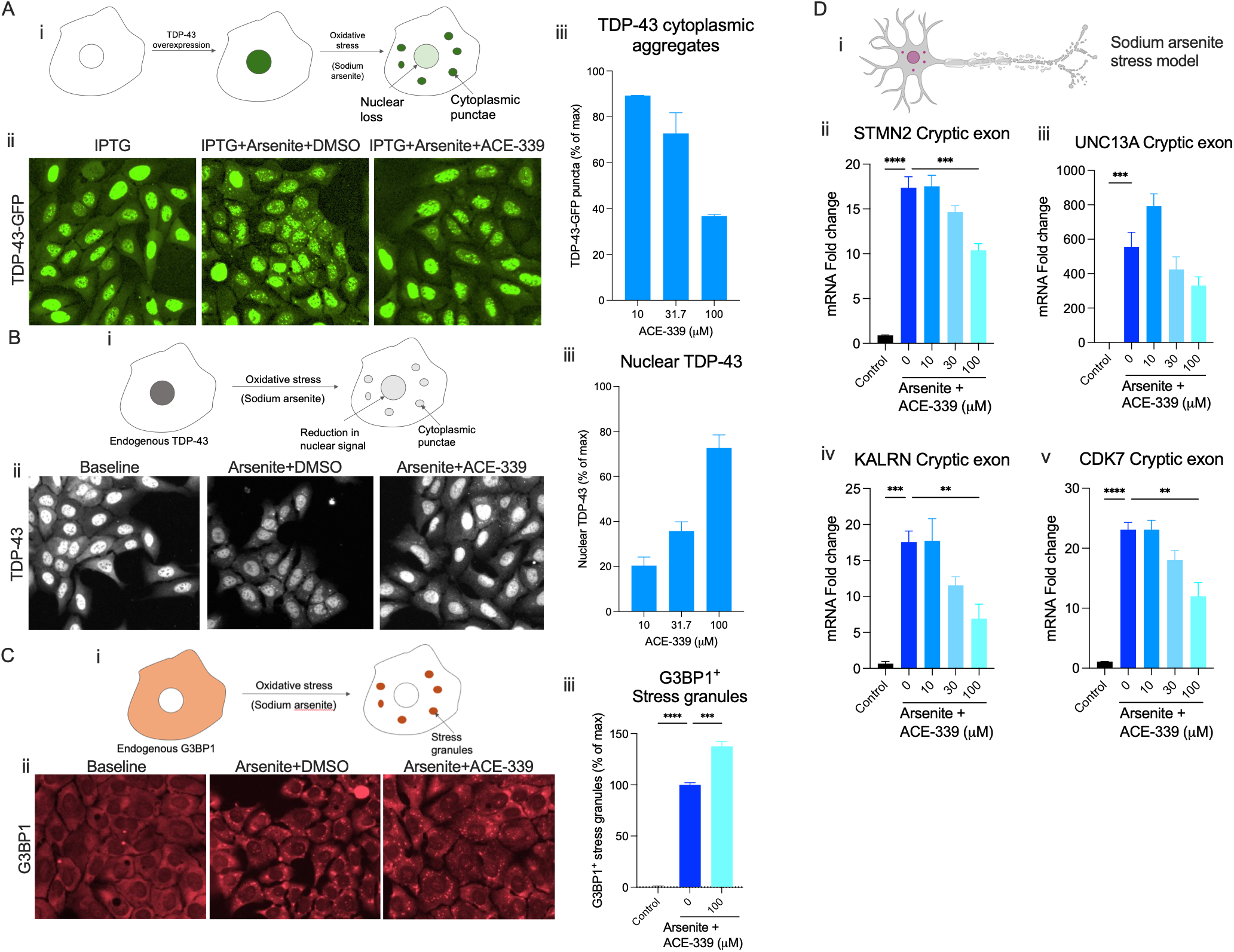
ACE-339 acts on cytoplasmic aggregated TDP-43 and rescues splicing function in oxidative stress model. (A) U2OS reporter cell model (i) overexpressing TDP-43-GFP (IPTG) was induced to form cytoplasmic puncta using 250μM sodium arsenite for 90 min in the presence of ACE-339 (24h pre-treatment). Representative images (ii) and quantification (iii) showing ACE-339 inhibits TDP-43-GFP cytoplasmic puncta. (B) U2OS cellular model for nuclear TDP-43 rescue (i) shows endogenous TDP-43 nuclear loss upon 250μM sodium arsenite treatment for 90 min at baseline conditions (non-overexpression, no IPTG). Representative images (at 100μM ACE-339) (ii) and quantification (iii) showing ACE-339 (24h pre-treatment) rescues nuclear TDP-43. (C) U2OS cellular model for stress granules (i) shows G3BP1+ stress granules upon 250μM sodium arsenite treatment for 90 min. Representative images (at 100µM ACE-339) (ii) and quantification (iii) showing ACE-339 (24h pre-treatment) does not affect stress granule formation. (D) ACE-339 rescues TDP-43 dysfunction dependent cryptic exons in indicated mRNAs in iPSC-derived motor neurons treated with sodium arsenite (250μM for 2h). Mean±SEM are shown. P<0.01**, P<0.001***, P<0.0001**** One-way ANOVA followed by Dunnett’s test. Image and quantification are representative of 2 independent runs with 2 biological replicates each.

In cells overexpressing TDP-43-GFP, we could not quantify a decrease in nuclear TDP-43 after arsenite stress induction, presumably due to interference from the excess TDP-43-GFP. We therefore used the U2OS-hTDP-43-GFP cells without overexpressing TDP-43-GFP, and observed a significant decrease in nuclear endogenous TDP-43 after arsenite stress (Figure 2Bi,ii). ACE-339 dose-dependently rescued endogenous TDP-43 nuclear localization (Figure 2Biii). Importantly, ACE-339 did not inhibit the formation of core G3BP1^+^ stress granules in cells stressed with sodium arsenite (Figure 2C), suggesting that ACE-339 specifically targets TDP-43 aggregate conformations without altering the normal cellular stress response. Interestingly, these G3BP1^+^ stress granules were somewhat smaller in size and greater in number in the presence of ACE-339, mimicking the observation in TDP-43 knockdown cells^38–40^. This minor effect on stress granules is consistent with a local reduction in TDP-43 within stress granules where it normally enhances their rate of fusion^38–40^. These data, paired with restored nuclear levels of TDP-43, suggest that ACE-339 specifically targets cytoplasmic TDP-43 conformations within puncta to support its localization in the nucleus.

When functionally localized within the nucleus, an important role of TDP-43 is to repress the inclusion of cryptic exons in mature mRNAs^41–47^. We reasoned that since arsenite stress induces TDP-43 cytoplasmic accumulation in puncta and reduced nuclear localization, it would result in less functional TDP-43 within the nucleus. We therefore quantified the abundance of TDP-43-regulated cryptic exons to measure the function of TDP-43 within the nucleus. We found that indeed, arsenite stress induction increased the amount of TDP-43-regulated cryptic exons in *STMN2*, *UNC13A*, *KALRN* and *CDK7* mRNAs. ACE-339 reduced the inclusion of all four of these TDP-43 dependent cryptic exons in stressed iPSC-derived human motor neurons (Figure 2D) as well as differentiated SH-SY5Y neuronal cells (Figure S5B). No significant changes were observed in total mRNA levels of *TARDBP*, *STMN2*, *UNC13A* or in cryptic exon inclusion in *HDGFL2*, *MYO18A*, *ACTL6B*, *POLDIP3*, *SYT7*, or *CAMK2B* upon arsenite stress. We note that high drug levels were required to rescue phenotypes in the arsenite induced stress model possibly due to the aggressiveness of the model, which results in massive displacement of TDP-43 from the nucleus to cytoplasmic puncta compared to both mouse models and human disease. In fact, these data are encouraging since they demonstrate the ability of ACE-339 to rescue pathology even under extreme stress. In support of this hypothesis, changes in cryptic exons in ALS patient derived iPSC-neurons are less than 2-fold^46,47^ as opposed to 15-fold or higher in the arsenite model (Figures 2D and S5B). We were unable to reproducibly detect TDP-43-dependent cryptic exons in other models like TDP-43 M337V mutant iPSC-derived motor neurons. Taken together, the recombinant protein and cellular data suggest that ACE-339 directly targets TDP-43 aggregation, thereby restoring physiologically folded, nuclear, functional TDP-43.

### ACE-339 displays high brain penetrance in wild type mice

In a pharmacokinetic study in wild type mice (Figure S6), ACE-339 was dosed orally, once daily, at 50, 150 and 400 mg/kg for 14 consecutive days. The drug was well tolerated at all doses. Plasma and brain drug levels measured on days 1 and 14 after dosing increased dose-dependently. On day 14, drug levels were lower compared to day 1 after dosing, possibly due to induction of liver metabolism by ACE-339. No significant dose-dependent serum biochemical changes were observed in plasma ALAT, ASAT, ALP, LDH, GGT, creatine kinase, creatinine, urea and total protein 24 hours after the last dose (not shown).

### ACE-339 reduces pathological TDP-43 and rescues pathology in iTDP43 mouse model

To test the target engagement of ACE-339 on pathological TDP-43 in target tissues and effects on downstream disease, we used a doxycycline (DOX) repressible model called iTDP43^A315T^ (or iTDP43) that overexpresses hTDP-43^A315T^ transgene under Thy1 promoter. The pathology in this model is driven by the human disease-associated A315T mutation which is known to induce insoluble, cytoplasmic, aggregated forms of TDP-43^48,49^. The iTDP43 mice (n=5 females+5 males/group) expressing the mutant transgene (DOX OFF) were treated with ACE-339 (75 and 150 mg/kg daily oral dosing) starting at 4 weeks and continued until 12 weeks of age (Figure 3A). ACE-339 was observed at high concentrations in the brains of these mice at the end of the study (Figure 3B). Additionally, ACE-339 reduced disease associated forms of TDP-43 in iTDP43 mice brains: detergent insoluble TDP-43, phosphorylated TDP-43, ubiquitinated TDP-43 and cytoplasmic TDP-43 (Figure 3C). Although some of these measures did not achieve statistical significance, a dose-dependent reduction in all of these biomarkers suggests meaningful effects of ACE-339 treatment. Consistent with a reduction in pathological TDP-43, ACE-339 reduced several disease phenotypes in the model. These include neuroinflammation, as indicated by a reduction in GFAP+ cells in the cortex and hippocampus of mice (Figure 3D), and improved motor function (pole test) and anxiety behavior (elevated plus maze, EPM) in iTDP43 mice. In the pole test, more ACE-339-treated mice were able to turn and descend the pole without falling, compared to vehicle-treated mice (Figure 3E and S7E). In EPM, ACE-339 treated iTDP43 mice spent less time in the open arm than vehicle-treated control mice (Figure S7F). ACE-339-treated mice also showed a reduction in time spent in the open arm of the EPM compared to vehicle-treated mice, suggesting a decrease in anxiety behavior (Figure 3E and S7G). ACE-339 did not affect hTDP43 transgene expression (Figure S7A), microgliosis (Figure S7B), rotarod motor behavior (Figure S7D), or the open field behavior test (Figure S7G). Neither the model nor ACE-339 treatment showed any effect on Ctip+ cortical layer V-VI neurons (Figure S7C) or animal body weight. No gender bias was observed in the mouse model or with ACE-339 treatment. These results indicate that ACE-339 engages and reduces pathological TDP-43 and downstream disease phenotypes in an iTDP43 mouse model.

**Figure 3.**
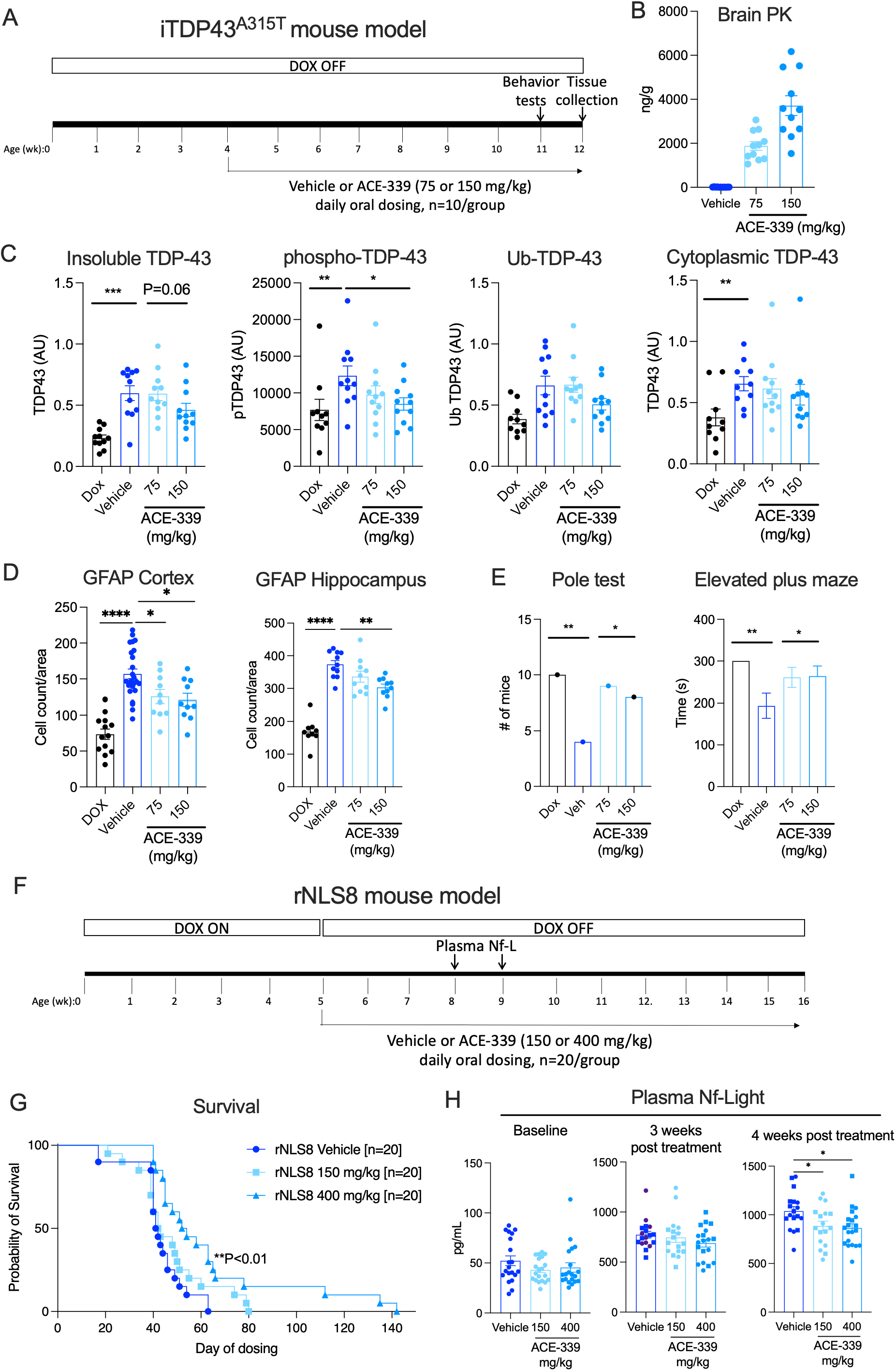
ACE-339 rescues disease phenotypes in two mouse models of TDP-43 pathology. (A) Schematic showing experimental design in iTDP43^A315T^ mouse model. Mice (n=10/group, 5 male and 5 female) were kept off doxycycline throughout the study and ACE-339 (75 and 150 mg/kg) was dosed orally, once daily from 4 weeks to 12 weeks of age. One group of mice was kept on DOX as non-transgenic-expression control. Behavior tests were done in the last week of dosing and tissues collected 2h after the last dose. (B) ACE-339 showed high dose dependent drug levels in the mice brain tissue (PK). (C) ACE-339 treatment reduced disease associated TDP-43: insoluble, phosphorylated, ubiquitinated and cytoplasmic TDP-43 in mouse brain tissue measured by western blotting. (D) ACE-339 rescues astrogliosis measured by GFAP immunostaining in mouse brain tissue. (E) ACE-339 treatment rescued motor (pole test, number of mice that completed the test) and anxiety behavior (Elevated Plus Maze, time spent exploring the maze) phenotypes. (F) Schematic showing experimental design in rNLS8 mouse model. Mice (n=20/group, 10 male and 10 female) were kept on DOX till 5 weeks of age. DOX was withdrawn and ACE-339 treatment (150 and 400 mg/kg, oral, once daily) began at 5 weeks and continued till the end of survival experiment. Plasma samples were collected from all mice for Nf-L measurement after 3 and 4 weeks of dosing (8 and 9 weeks of age respectively). (G) ACE-339 significantly rescued survival at both doses in rNLS8 mice. P<0.01** Mantel-Cox test. (H) Plasma Nf-light measurement at baseline (pre-treatment), 3 week and 4 weeks after dosing. ACE-339 significantly rescued Nf-L after 4 weeks of dosing. Mean±SEM are shown. P<0.05*, P<0.01**, P<0.001***, P<0.0001**** One-way ANOVA followed by Dunnett’s test.

### ACE-339 rescues survival and Nf-L in rNLS8 mouse model

We also assessed the efficacy of ACE-339 in another mouse model displaying additional human disease relevant biomarkers. The rNLS8 mouse model overexpresses hTDP-43 lacking the nuclear localization signal (hTDP43-ΔNLS) driven by a DOX repressible NEFH promoter to drive disease. Transgenic hTDP43-ΔNLS remains in the cytoplasm where it recruits endogenous mouse TDP-43 thereby driving pathology^50,51^. We hypothesized that ACE-339 would target cytoplasmic, aggregated hTDP43-ΔNLS and that, although this protein itself cannot localize to the nucleus, endogenous mouse TDP-43 would re-localize to the nucleus and rescue pathology. Importantly, rNLS8 mice exhibit human disease relevant biomarkers upon DOX withdrawal such as short survival periods and high plasma neurofilament-light chain (Nf-L). These phenotypes are absent in iTDP43 mice.

rNLS8 mice (n=10 females+10 males/group) were treated with ACE-339 (150 and 400mg/kg daily oral dosing) starting when DOX was withdrawn at 5 weeks of age (Figure 3F). Remarkably, ACE-339 dose-dependently delayed disease onset and improved survival in rNLS8 mice (Figure 3G). While all vehicle treated mice died by day 63, mice in the low dose treatment group survived up to 80 days. Mice in the high dose group survived up to 145 days after DOX withdrawal, with a median survival rescue of 11.5 days (∼ 28%) (P<0.01).

Additionally, both doses reduced plasma Nf-L after 4 weeks of ACE-339 treatment (Figure 3H). No effect on Nf-L was observed at 3 weeks of ACE-339 treatment, likely because the most severely affected vehicle-treated mice died before this time point. No significant gender bias was observed with ACE-339 treatment (Figure S8A, B) except that the 2 mice that lived beyond 126 days in the high dose group were both females. Additionally, ACE-339 treatment reduced body weight loss in these mice (Figure S8C). Drug measurement in a satellite group of rNLS8 mice showed brain penetrance, albeit lower than that observed in wild type mice, possibly due to the mouse strain difference (Figure S8D). Importantly, ACE-339 reduced the disease associated insoluble form of TDP-43 in rNLS8 mice demonstrating target engagement in the brain (Figure S8E).

To evaluate the downstream transcriptomic impact of ACE-339, we performed RNA-seq on rNLS8 mouse brains from this study. ACE-339 treatment induced a distinct transcriptome-wide isoform expression shift, significantly depleting select isoforms in genes associated with cell death and neuroinflammatory pathways while increasing select isoforms in genes involved in neurogenesis and neurodevelopmental pathways (Figure S9). These data further support that target engagement by ACE-339 with pathological TDP-43 successfully suppresses neurotoxic inflammation and promotes neuronal repair *in vivo*.

### TDP-43 target load in human ALS and mouse model tissues

We reasoned that due to the distinct ACE-339 mechanism of action and the overexpression of TDP-43 in the mouse models, a direct translation of mouse to human doses using traditional allometric scaling would not be appropriate and should instead be considered relative to the amount of TDP-43. To address this, we quantified TDP-43 protein in brain tissues from post-mortem ALS patients, iTDP43 and rNLS8 mouse models. Total TDP-43 concentration in ALS brain tissues was about 20 and 200-fold less than iTDP43^A315T^ and rNLS8 mouse brain tissues respectively (Figure S10). These data demonstrate high TDP-43 overexpression in these mouse models. The ACE-339 doses selected for the respective mouse model were based on these high TDP-43 concentrations in the brain tissues. Despite the high target load, ACE-339 showed activity in both of these severe models of TDP-43 pathology.

### Molecular dynamics simulations reveal TDP-43 hexameric structure and its association with ACE-339

To simulate the assembly of TDP-43 (307-319) into a beta-sheet rich hexamer and interrogate how this system interacts with ACE-339, a molecular dynamics (MD) computer simulation methodology was deployed that forms a component part of Acelot’s RESOLUTE platform. These MD simulations suggest that ACE-339 engages with TDP-43 oligomers through cryptic pocket engagements. Firstly, in order to create a representative hexameric structure (chosen based on the previously hexameric oligomers observed via mass spectrometry^25^) to perform atomic simulations, replica exchange molecular dynamics (REMD) methodology was deployed starting with a molecular geometry consisting of 6 randomly posed misfolded TDP-43 peptides. During the course of multiple simulations these peptide sequences began to assemble into beta-barrel hexamers (Figure 4A) with prominent cross-sequence interactions between residues ^311^M, ^313^F, and ^316^F (Figure S11A). Analysis of these molecular simulation trajectories highlighted several prominent architectural arrangements of oligomer structural elements, with one in particular with a prevalence of 7.8%, which shall be referred to here as the ’pre-formed TDP-43 hexamer’. We used this pre-formed hexameric geometry as a starting point for additional unbiased MD simulations including ACE-339 in order to study its binding characteristics and what intermolecular interactions are engaged with TDP-43 hexamers to shed light on how ACE-339 experimentally disaggregates TDP-43. Using a cluster analysis of poses from these simulations, we found that ACE-339 displays 4 unique putative binding sites in this pre-formed TDP-43 hexamer (Figure 4B), suggesting dynamic engagement of a potential cryptic pocket. To study this cryptic pocket binding hypothesis in more detail, additional REMD simulation trajectories were performed, where ACE-339 showed greater binding interaction contacts around the 311M, 313F, and 316F amino acid side chains (Figure S11B).

**Figure 4.**
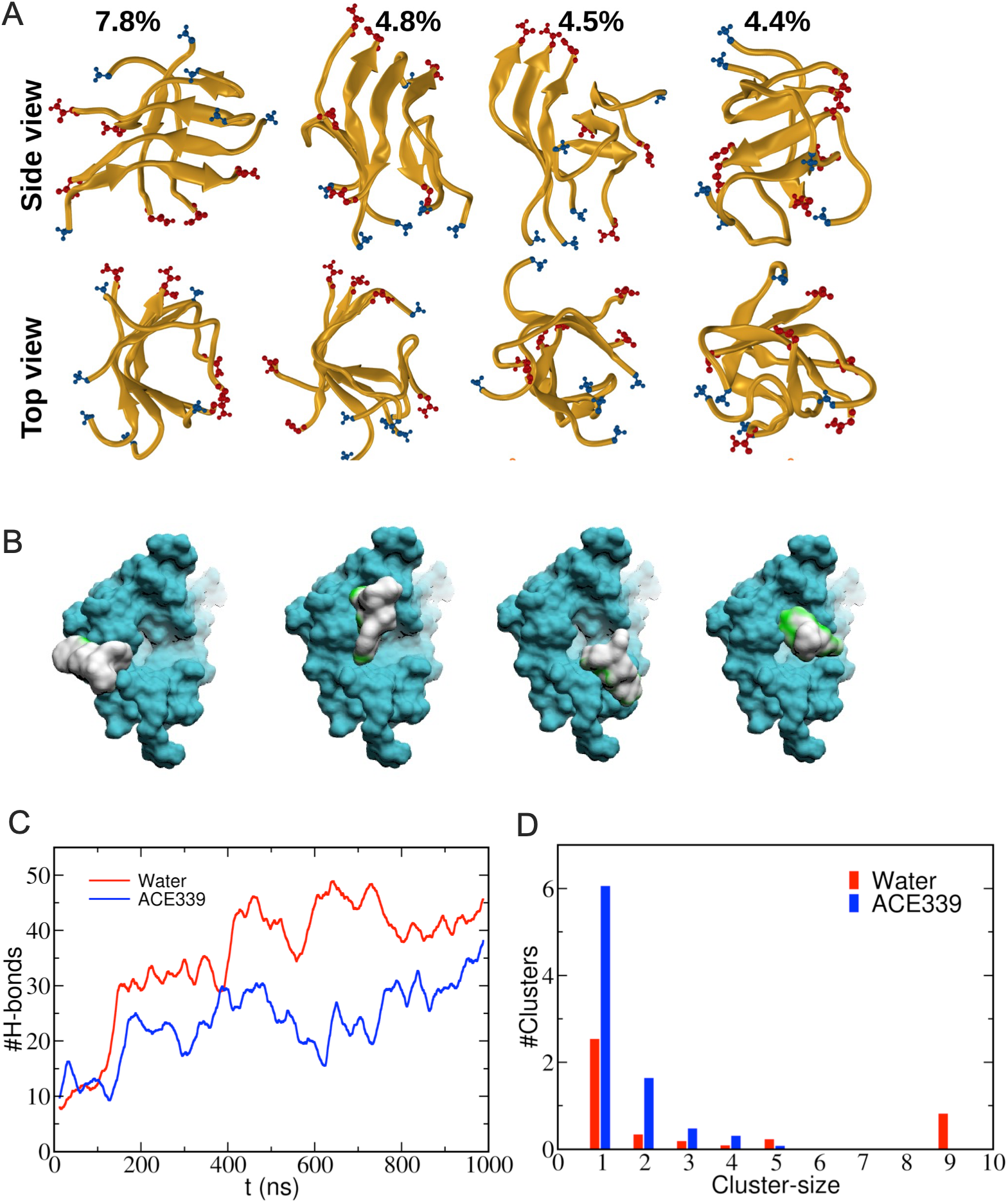
ACE-339 binds to hexameric TDP-43 in molecular dynamics simulations. Replica exchange molecular dynamics (REMD) simulations were conducted by inserting six TDP-43 peptides at random positions in a simulation box, solvated in water (100 mM NaCl buffer), 1100 ns long trajectories were obtained with 64 replicas with the final 300 ns of trajectory time at 300 K being used for data analyses. (A) The most probable conformations (relative percentages shown) of TDP-43 (307-319) hexamers in aqueous solution. (B) ACE-339 (white with green areas of high engagement) binding interactions with TDP-43 (307-319) hexamer (turquoise), in 4 positions based on cluster analysis superimposed on one TDP-43 simulation snapshot to clearly display all 4 positions. (C-D) Twelve TDP-43 (307-319) molecules were randomly inserted into a large simulation box with a linear dimension of 25 nm, resulting in a peptide concentration of approximately 1mM. These simulations were carried out for 1 microsecond each. (C) ACE-339 reduced the total number of intrapeptide and interpeptide hydrogen bonds in the TDP-43 oligomers. (D) ACE-339 reduces the TDP-43 peptide cluster size (number of peptides). A cutoff distance of 0.6nm between the backbone atoms averaged over 500-1000ns was used for identifying the clusters.

Due to the limitations of REMD, which at higher peptide concentrations resulted in discontinuous trajectories leading to a loss of information during hexamer formation, additional unbiased MD simulations were conducted at 300K. Twelve TDP-43 (307-319) molecules were randomly inserted into a large simulation box with a linear dimension of 25 nm, resulting in a peptide concentration of approximately 1mM. These simulations were carried out for 1 microsecond each. The time evolution of peptide aggregation was analyzed by monitoring the total inter-peptide and intra-peptide hydrogen bonds over the course of the simulation. Inclusion of ACE-339 led to a significant reduction of the total number of hydrogen bonds formed between TDP-43 peptides (Figure 4C). Furthermore, while TDP-43 formed large oligomers with aggregate sizes extending up to 9-mers in this timeframe, it was unable to do so in the presence of ACE-339 (Figure 4D). Together, these *in silico* findings add support to the ability of ACE-339 to alter the equilibrium of the TDP-43 aggregation pathway, pushing it favorably towards the restoration of functional TDP-43.

## Discussion

The association of TDP-43 with the pathogenesis of neurodegenerative diseases like ALS and FTLD-TDP has been extensively described, but the causes of TDP-43 aggregation are still debated. Mechanisms like stress granule formation via the integrated stress response pathway and nuclear transport defects have been implicated in TDP-43 dysfunction in diseases^2,20^. Both TDP-43 aggregation (gain-of-function) and nuclear TDP-43 clearance (loss-of-function) are considered disease drivers. We developed small molecules that target TDP-43 aggregation with the hypothesis that in disassembling pathological aggregated TDP-43, more functional TDP-43 would become available to restore loss-of-function phenotypes such as TDP-43 nuclear loss, downstream splicing functions, and neuronal cell death, with the aim to ameliorate disease. We show that ACE-339 directly targets aggregated (misfolded, insoluble) TDP-43 and in doing so, it also rescues TDP-43 localization, downstream functional phenotypes, and ameliorates *in vivo* disease models. These small molecules are orally bioavailable and highly cell- and brain-penetrant, which is crucial for targeting intracellular proteins in neurodegenerative diseases.

To elucidate how ACE-339 clears TDP-43 pathology *in vivo*, we evaluated its direct structural and molecular binding dynamics. Microscale thermophoresis and target-engagement assays suggest that ACE-339 binds TDP-43 in a strictly conformation-specific manner: it interacts directly with aggregated TDP-43 but does not engage with natively folded TDP-43 monomers. To explore the physical basis of this selectivity, we interrogated the binding interface using molecular dynamics simulations. These simulations suggest that ACE-339 fits directly into the highly aggregation-prone core of TDP-43 (residues 307-319). Rather than binding a single static surface pocket, ACE-339 appears to use multiple distinct chemical groups to dynamically engage the tightly packed interface between the aggregated peptide chains. By wedging into this interface, ACE-339 disrupts critical inter-peptide hydrogen bonds, effectively prying the aggregated beta-sheets apart.

Once this localized aggregate architecture is dismantled and TDP-43 reverts to a monomer, the pathological binding interface is eliminated. Fractionation of disaggregated TDP-43 confirmed that upon aggregate dissolution, ACE-339 is subsequently released back into the free fraction. Taken together, these data are consistent with a structural mechanism in which ACE-339 selectively targets and physically dismantles the TDP-43 aggregate core. Because ACE-339 loses its binding site upon target dissociation, sub-stoichiometric concentrations can iteratively engage multiple aggregates. We propose that this dynamic equilibrium enhances the clearance of insoluble pathology observed in the mouse models used in this study.

Crucially, by disrupting the aggregate, ACE-339 releases native TDP-43 monomers that are competent to refold and translocate back into the nucleus. This therapeutic disaggregation not only clears toxic cytoplasmic inclusions but actively restores the functional pool of TDP-43 required for normal cellular regulation, as supported by the functional rescue observed in cellular models and the reversal of in vivo disease phenotypes, including prolonged survival, reduced Nf-L release, and reduced neuroinflammation.

While investigating the mechanism of ACE-339 activity, we generated experimental support for the under-appreciated notion of TDP-43 oligomer toxicity^23–26,52,53^. This idea is derived from amyloid beta research that shows smaller oligomers of amyloid beta and tau, rather than larger aggregates/plaques, lead to toxicity in Alzheimer’s disease. The large aggregates/plaques are in fact considered protective as they can sequester the toxic oligomers. Similarly, alpha-synuclein oligomers are thought to be toxic in Parkinson’s disease. We demonstrated that TDP-43 oligomers can cause synaptic defects and ACE-339 targets these oligomers to rescue synaptic function. Since ACE-339 also disrupted mature TDP-43 fibrils, we hypothesize that the large aggregates are in a dynamic equilibrium with oligomers, which is supported by our observations that extending the incubation of TDP-43 oligomers forms fibrils, and that ACE-339 acts on the oligomers to further push the equilibrium toward functional forms of TDP-43.

In this study, we utilized several overexpression and stress models to assess ACE-339 activity. These models were chosen for the robustness of the phenotypes induced. Without such aggressive manipulations, TDP-43 aggregation dependent phenotypes cannot be clearly observed in cellular and mouse models. ACE-339 efficacy in such severe models with high TDP-43 aggregate burden suggests that ACE-339 may be effective in the milder, more physiological disease conditions seen in human disease^47^ at lower doses.

Familial mutations in TDP-43 are associated with accelerated disease pathology in patients, however the majority of patients have no known mutations. TDP-43 (A315T) is one such mutation associated with ALS/FTD. Patients with this mutation were shown to progressively develop lower motor neuron degeneration with respiratory failure^54,55^. ACE-339 showed activity in the mouse model with this TDP-43 (A315T) mutation suggesting that ACE-339 can be beneficial for the genetic ALS subpopulation as well. Since ACE-339 interacts specifically with the aggregated form of TDP-43 and not the native conformation, despite sharing the same primary amino acid sequence, we hypothesize that ACE-339 may have activity on other TDP-43 mutants as well. While large, hyperphosphorylated TDP-43 cytoplasmic inclusions represent the classical diagnostic hallmark of ALS, neuropathological staging suggests that TDP-43 aggregation may occur earlier as evidenced by TDP-43 appearance as granular cytoplasmic ‘dust-like particles’ that precede mature compact inclusions^56–58^. Furthermore, recent detection of pathological TDP-43 using conformation-selective RNA aptamers suggests that early pathological TDP-43 self-assembly is far more pervasive across human ALS CNS tissue than indicated by classical inclusion histology, directly coinciding with the cryptic splicing loss-of-function pathology^59^. We designed TDP-43 disaggregators that may function as disease modifiers by not only targeting TDP-43 aggregation, but also by restoring its nuclear, functional forms. The ability of ACE-339 to disrupt distinctly derived versions of TDP-43 aggregates and oligomers, including four different recombinant forms, two different cellular-induced forms, and two different induced forms *in vivo*, suggests that ACE-339 may interfere with pathological TDP-43 in a wide variety of contexts or conformations.

The ability to disrupt multiple aggregate species translated into meaningful benefit *in vivo*. In disease mice, ACE-339 significantly extended lifespan and reduced Nf-L, two phenotypes relevant to human ALS. Nf-L reduction in particular has become a key marker of therapeutic response in the clinic: the FDA-approved SOD1 ASO, Tofersen (Qalsody ®), improved survival in the open-label extension and showed a 50-60% decrease in Nf-L over 28 weeks of treatment in familial patients with SOD1 mutation. However, SOD1 mutations account for only ∼2% of ALS cases, whereas TDP-43 pathology is present in ∼97%. Other FDA-approved drugs for ALS (Riluzole and Edaravone) largely relieve symptoms without targeting the underlying disease pathology and many drugs targeting other pathways have not shown efficacy in ALS and FTLD clinical trials. Other drugs directly targeting TDP-43 are under development and have different modalities (antibodies, DNA/RNA-based therapies)^60^. Small molecules like ACE-339 are particularly promising because they can act inside cells on specific disease associated forms of TDP-43. That could benefit not only sporadic and familial ALS and FTLD, but possibly other TDP-43 proteinopathies too, such as inclusion body myositis, multiple systems proteinopathy (MSP), limbic-predominant age-related TDP-43 encephalopathy (LATE), facial onset sensory and motor neuronopathy (FOSMN) and diseases like Alzheimer’s and Parkinson’s which may have TDP-43 co-pathology.

The *in silico* identification of ACE-339 marks a significant advancement in the field of ALS drug discovery as a compound discovered solely via machine learning. The comprehensive characterization of ACE-339’s drug-like properties (not shown here) not only underscores its potential therapeutic value but also highlights the increasing role of Artificial Intelligence in accelerating the drug development processes. Our approach illustrates how machine learning and computational chemistry approaches like the RESOLUTE platform can streamline the discovery of novel compounds, ultimately reducing the time and cost associated with bringing effective drugs to patients. Given the success of the RESOLUTE platform targeting this historically difficult protein aggregate target, the platform is currently being expanded to explore a variety of allosteric or cryptic pocket and protein-protein interactions drug discovery challenges.

## Methods

### Ethics Statement

Post-mortem human frontal cortex tissues from ALS patients (n = 3) were obtained from the Institute for Brain Aging and Dementia at the University of California, Irvine, with approval from the Institutional Ethics Committee, and post-mortem motor cortex tissues from sporadic ALS patients (n = 14) and non-neurological controls (n = 8) were obtained from Target ALS biobanks at Georgetown University and the University of Edinburgh under their respective institutional ethical approvals. All animal procedures were conducted in accordance with institutional and national animal welfare guidelines and were approved by the Macquarie University Animal Ethics Committee (*iTDP-43^A^*^315^*^T^* study), the PsychoGenics Inc. Institutional Animal Care and Use Committee (*rNLS8* study), and the Enamine/Bienta Institutional Animal Care and Use Committee (wild-type mouse pharmacokinetic study).

### TDP-43 peptide synthesis and ThT assay

The effect of compound on fibril formation by TDP-43 (307-319) peptide was evaluated using a thioflavin T (ThT) fluorescence assay. Monomeric, lyophilized TDP-43 (307-319) (synthesized at Genscript) was reconstituted in HFIP to generate a 3mM stock solution and stored in -80°C till use. This stock solution was diluted with 100 mM ammonium acetate (Sigma-Aldrich, 09691-100ML) to attain a working concentration of 200 μM on ice. Compounds were dissolved in 100% dimethyl sulfoxide (DMSO) (MilliporeSigma, D2650) to make a 10 mM stock and stored at -20°C. A reaction mixture, containing 30 µM ThT (VWR, 103011-384), 50 µM TDP-43 (307-319), 2.5% HFIP, and 100 mM ammonium acetate was prepared and maintained on ice. The reaction mixture was then dispensed into multiple wells of a 96-well half-area, low-binding polyethylene non-treated plate (Corning, 3880) with a clear bottom at a volume of 98 μL per well. To each respective well, 2 μL of 10 mM compound stock was added in duplicates at 0h for coincubation and at 6h for disaggregation assay. The vehicle control consisted of 30 µM ThT, 50 µM TDP-43 peptide, 2.5% HFIP and 2% DMSO in 100 mM ammonium acetate, while the blank control comprised 30 µM ThT, 2.5% HFIP, and 2% DMSO in 100 mM ammonium acetate. Plates were sealed and placed in a plate reader (BioTek Synergy Neo2) at 37°C without agitation. ThT fluorescence was measured through the bottom of the plate using 440- and 490-nm excitation and emission filters for 24-25 hours at 0.5-hour intervals, scanning each well at 5 points. The initial screening assay was performed at a 1:4 stoichiometric ratio with 50µM peptide and 200µM compound.

To determine the IC_50_, 10mM stock of ACE-339 was serially diluted, 5μL of compound stocks were added to the 50µM TDP-43 (307-319) mix described above in duplicates. For data analysis, the RFU at 24.5 hours was used to calculate the percent of aggregation for all concentrations, Blank controls were first subtracted from all groups, with vehicle considered 100% aggregated. All compound groups were normalized to vehicle control to calculate the percentage aggregation, followed by 4PL curve fitting and IC_50_ calculation using GraphPad Prism.

### TDP-43 Aggregation and Dot Blot Analysis

Recombinant full-length TDP-43 (Genscript, supplied in 8M urea, 50mM Tris buffer) was introduced into native buffer (1X PBS, 5mM DTT) by fast dilution to a final concentration of 5μM and allowed to aggregate for one week at room temperature under quiescent conditions. Following aggregation, the sample was mixed by pipetting and diluted to 100nM in native buffer and incubated with either ACE-339 (500nM) or equivalent DMSO for 24 hours at room temperature. Samples were then centrifuged at 21,100 × g for 1 hour at room temperature to separate soluble and insoluble fractions, and all fractions were treated with 2% SDS at 90°C for 10 minutes prior to blotting. Dot blots were performed by spotting 5μL of each fraction onto nitrocellulose membranes, blocked with Odyssey Blocking Buffer, and probed with anti-TDP-43 antibody (Proteintech, 10782-2-AP) and goat anti-rabbit AzureSpectra 800 secondary antibody. Membranes were imaged on a Li-Cor Odyssey CLx and dot intensities quantified using Li-Cor Image Studio software. Data were analyzed using GraphPad Prism.

### Human disease tissue derived TDP-43 oligomer assays

#### Human brain tissue harvesting

As described previously^61^. Frontal cortex brain tissues from ALS patients (n = 3) were received as frozen blocks from the Institute for Brain Aging and Dementia at UC Irvine following approval by the Institutional Ethics Committee. Brain tissues were homogenized in 1× PBS mixed with a protease inhibitor cocktail (Roche, #11836145001) and phosphatase inhibitor (Sigma, #04906837001) at 1:3 (w/v) dilution of brain:PBS. Samples were then centrifuged at 10,000 xg for 10 min at 4 °C. The supernatants (PBS-soluble fractions) were aliquoted, snap-frozen, and stored at −80 °C until use. The pellets were re-suspended in Sarkosyl solution at a final concentration of 1% with stirring for 1 h at RT and then centrifuged at 100,000 xg for 30 min at 4 °C. Sarkosyl-soluble fractions (PBS-insoluble fractions) were aliquoted, snap-frozen, and stored at −80 °C until use. Characteristics of the human brain tissues are summarized in Table S1.

#### Brain-derived TDP-43 oligomer isolation

The PBS-soluble fractions of ALS brain homogenates were immunoprecipitated with TDP43-Oligo01 antibody as described previously^26^. The Pierce™ Co-Immunoprecipitation Kit (Thermo Scientific, #26149) was used following the manufacturer’s guidelines. Briefly, amine-reactive resin was coupled with the TDP-43-Oligo01 antibody, which is suitable for IP, followed by incubation with brain homogenates (PBS-soluble fractions). Bound proteins were eluted in 0.1 M glycine (pH 2.8), and the final pH was adjusted to 7.0 by adding 1 M Tris-HCl (pH 8). Isolated fractions were subjected to buffer exchange and finally collected in sterile PBS followed by IB characterization. The total protein concentrations were measured with bicinchoninic acid protein assays (Micro BCA Kit, Pierce).

#### Full length TDP-43 (FL TDP-43) expression, purification

Recombinant FL TDP-43 was purified using the method previously described^62^. *Escherichia coli* BL21 (DE3) competent cells (New England Biolabs) were used to express TDP-43 FL protein using the introduced plasmid (pCOLD TDP43 FL). Cryopreserved transformed *E.coli* cells were grown in Luria-Bertani (LB) broth, supplemented with 1mg/ml ampicillin. The cells were grown at 37°C with shaking at 220 rpm till the O.D_600_ reached a value of 0.6-0.8, following which protein over-expression was induced using 1M IPTG (Isopropyl β-D-1-thiogalactopyranoside) and incubating at 16 °C, while shaking at 150 rpm for ∼16 h. Cells were harvested by centrifugation at 5000 rpm for 10 mins at 4°C. The cell pellets were re-suspended in 25ml lysis buffer (40 mM Hepes-KOH, 500mM KCl, 20mM MgCl2, 10% glycerol, 20 mM β-mercaptoethanol, pH 7.4) supplemented with one cOmplete ™, EDTA-free Protease Inhibitor Cocktail (Sigma-Aldrich) and 150 mM PMSF. The cells were lysed by sonication; 45 secs burst, followed by 1 min rest on ice, which was repeated ten times. Lysates were centrifuged at 12000 rpm for 15min at 4°C. and supernatants collected, incubated with HisPur™ Ni-NTA Resin (Thermo) for a minimum of 30 mins at 4°C. The mixture was then packed into a Kontes® flex-column. The column was washed with 2 bed volumes of wash buffer (40 mM Hepes-KOH, 500mM KCl, 20mM MgCl2, 10% glycerol, 20 mM β-mercaptoethanol pH 7.4) and eluted in elution buffer (50mM Tris HCl, 150mM NaCl, 0.5M arginine and 10% glycerol and 250 mM Imidazole pH 7.4). TDP-43 purified with this method was used for amplifying oligomers, see supplemental methods for purification of FL TDP-43 used in ThT and MST assays.

#### Amplification of seeded TDP-43 oligomers and fibrils from brain derived oligomers

Oligomers of recombinant FL TDP-43 were amplified by seeding with TDP-43 oligomer immune-precipitates (IP) from ALS brain tissues as previously described^26,52^. The IP and TDP-43 monomer mixture was made at a ratio of 1:100 (molar ratio), left for gentle agitation at room temperature for 48hrs. After seeding and amplification, aliquots were taken and immediately used for immunoblotting (IB) using A11-19 and TDP-43 oligomer specific antibodies as well as atomic force microscopy analysis for oligomer morphology and quality control. Fibrils were formed at a similar experimental condition used for oligomers described above but need extended time. The samples were placed on rotation at 10-15 rpm for 5 days at 37 °C.

#### Immunochemistry of oligomers (Dot blot)

To detect FL TDP-43 oligomer before and after the treatment with ACE-339, dot blot assay was used as described in^63^. 1.5μl sample was applied onto a 0.45μm nitrocellulose membrane (Cytiva Amersham™ Protran™). The membrane was blocked with 10% nonfat milk in TBS with low tween (TBST) for 1.5h at RT. After that, membrane was probed with A11-19 antibody diluted in 5% nonfat milk and incubated for 1h at RT. The primary antibody was removed washing the membrane three-times with TBST for 10 min and incubated for 1h at RT with anti-rabbit secondary antibody diluted in TBST (1:20000). ECL developer solution (Advansta WesternBright^TM^ ECL) was applied for 2 min before signal detection on X-ray film.

#### Native-PAGE and Silver Stain

Native-PAGE 3-12% Bis-tris (Invitrogen) precast gel was used for native-PAGE analysis. The comb was removed from the gel and inserted into the XCell SureLOck Mini-cell gel running tank. The wells were rinsed three times with 1X NativePAGE Dark Blue Cathode Buffer. The white tape near the bottom of the gel cassette was removed, and the gel wells were filled with 1X NativePAGE dark blue cathode buffer. Two micrograms of each sample were prepared in Native-PAGE 1X loading dye (Invitrogen) and loaded into the wells. The inner chamber was filled with 200 mL 1X NativePAGE dark blue cathode buffer and the outer chamber with 550 mL 1X NativePAGE Anode Buffer. The gel was run at 100 V until the dye front reached one-third of the gel length, then the dark blue cathode buffer in the inner chamber was replaced with light blue cathode buffer, and the gel was run at 120 V for 2 hours. The gel was removed and destained with a solution of 40% methanol, 50% water, and 10% acetic acid, then subjected to silver staining for protein band visualization. The Thermo Scientific™ Pierce™ Silver Stain Kit was used for rapid and ultrasensitive protein detection in polyacrylamide gels. The gel was washed twice for 5 minutes each in ultrapure water, fixed twice for 15 minutes each in 30% ethanol and 10% acetic acid solution, again washed twice for 5 minutes in 10% ethanol, then twice for 5 minutes in ultrapure water. The gel was sensitized for 1 minute in sensitizer solution (50 μL Sensitizer with 25 mL water), then washed twice for 1 minute with water. The gel was stained for 30 minutes in stain working solution (0.5 mL enhancer with 25 mL stain), washed twice for 20 seconds with ultrapure water, and developed for 2-3 minutes in developer working solution (0.5 mL enhancer with 25 mL developer) until bands appeared. Development was stopped with 5% acetic acid for 10 minutes, and images of the gel were taken.

#### Atomic force microscopy (AFM)

AFM images of oligomer of FL TDP-43 alone (10 µM) and treated with ACE-339 (0 µM, 100µM and 200µM) were obtained using a non-contacting tapping method with Bruker Multimode 8 AFM. A 12mm mica fixed on a 15 mm metallic disc was freshly cleaved to obtain a uniform surface. 10µL of 1:10 dilution sample (0.03-0.05 µg/µL) in PBS was applied on mica and allowed to adsorb overnight (∼16h at RT). Mica was then repeatedly washed with molecular grade water and air dried. Images were taken of at least three (max of five) different fields on the mica surface. Finally, images were analyzed using NanoScope Analysis v1.20rl AFM data processing software and data was plotted using GraphPad Prism.

For AFM imaging of fibrils, 2 μl of fibril sample was diluted 10-fold in molecular grade water, applied on freshly cleaved mica plate and dried air overnight. The surface was then washed 3 times with molecular grade water and air dried. Before imaging, scan parameters were set as scan size of 10µm, 5µm and 2 µm, aspect ratio of 1.0. Keep on ScanAsyst auto control and data type as “height” to scan image. Images were taken with a Multimode 8 AFM machine (Veeco, CA) using a non-contact tapping method (ScanAsyst-Air). AFM analyses were performed using the particles analysis tool of the NanoScope Analysis v1.20rl AFM data processing software. The reported height maxima and periodicity of fibrils are based on measurements using 4 randomly selected fibrils.

#### Electrophysiology

Acute brain slices were prepared as previously described^64^. Mice were deeply anaesthetized, quickly decapitated, and the brain was extracted. Transverse brain sections of 375 μm containing Schaffer collateral synapses were generated using the Compresstome VF-300 (Precisionary Instruments, Greenville, NC, USA). Slices were allowed to recover in NMDG-HEPES cutting solution at 32°C for 30min, during which a Na^+^ spike-in protocol was performed. The slices were then transferred to carbogen bubbling HEPES-holding solution at room temperature for the rest of the experiment time. Slices were treated with oligomers after 2 hours of recovery in HEPES-holding solution. For treatment, the slices were transferred to separate chambers and incubated with 100nM of each oligomeric sample (with 0, 1µM or 2µM ACE-339) diluted in carbogen bubbling HEPES-holding solution at room temperature for 50min before recording. After treatment, the slices were briefly (5 min) washed by placing them in oligomer-free HEPES-holding solution before placing them on the recording stage. Two slices at a time were transferred to the recording chamber and perfused with carbogen bubbling room temperature normal artificial cerebrospinal fluid (aCSF) at a rate of approximately 3 ml/min for recordings. Recording electrodes were pulled from borosilicate glass capillaries using a horizontal P-97 Flaming/Brown micropipette puller (Sutter Instruments, Novato, CA). Evoked field excitatory post-synaptic potentials (fEPSPs) in the CA1 were obtained by stimulating Schaffer collateral. A stable baseline was obtained by delivering single-pulse stimulation at 20-s interstimulus intervals. Long-term potentiation (LTP) was induced by exposing the slices to high-frequency stimulation (HFS; 3 × 100 Hz, 20s intervals) using Digidata 1550B (Molecular Devices, Sunnyvale, CA, USA), and measured using an Axon MultiClamp 700B differential amplifier (Molecular Devices). Clampex 10.6 software (Molecular Devices) was used. All data are represented as percent change from the initial average baseline fEPSP slope, which was defined as the average slope obtained for 10 min before HFS.

### Cellular TDP-43 assays

#### U2OS hTDP-43-GFP arsenite-induced puncta

U2OS reporter cells stably expressing a two-part IPTG-inducible system for expression of human full-length TDP-43 C-terminally tagged with GFP were purchased from Innoprot (Spain) and cultured in DMEM/F-12 media with HEPES and 10% FBS in 0.17 um thin cyclic olefin-bottom 96-well imaging plates. One day after plating, cells were treated with 5mM IPTG and DMSO or ACE-339 for 20-24 hours. Sodium arsenite (or water control) was spiked into the culture media to 250µM for 90 minutes and cells were fixed with 4% paraformaldehyde and stained with 10µM Hoechst 33342 for 1 hour prior to imaging. TDP-43-GFP puncta were identified in the cellular region excluding the nucleus using IN Carta image analysis software. The initial screening assay was performed with 100µM compound.

#### U2OS stress-free hTDP-43-GFP puncta

Cells were prepared as above. One day after plating, cells were treated with 5mM IPTG and DMSO or ACE-339 for 20-24 hours. Cells were fixed with 4% paraformaldehyde and stained with 10µM Hoechst 33342 for 1 hour prior to imaging. TDP-43-GFP puncta were identified in the cellular region excluding the nucleus using IN Carta image analysis software.

#### U2OS arsenite-induced TDP-43 nuclear depletion

U2OS reporter cells were cultured and seeded as above, and maintained without IPTG induction. One day after seeding, cells were treated with DMSO or ACE-339 for 20-24 hours. Sodium arsenite (or water control) was spiked into the culture media to 250µM for 90 minutes and cells were fixed with 4% paraformaldehyde. Cells were permeabilized with cold 0.2% TX-100/PBS for 5 minutes, blocked with 10% normal donkey serum (NDS) (Jackson ImmunoResearch Laboratories Inc) for 1 hour, incubated overnight with 1:200 rabbit anti-TDP-43 (Proteintech 10782-2-AP; N-terminal antibody), incubated with 1:250 Alexa 647-conjugated donkey anti-rabbit secondary antibodies and 10µM Hoechst 33342 for 30 minutes prior to imaging. The total intensity of TDP-43 was measured within the nuclear region stained by Hoechst.

#### U2OS arsenite-induced G3BP+ stress granules

U2OS reporter cells were cultured, seeded, and treated as above, and concomitant with staining for endogenous TDP-43, cells were incubated with mouse anti-G3BP-1 (ab56574, Abcam, 1:3200) with Rhodamine Red X-conjugated secondary antibodies.

#### Microscopy and image analysis

Fixed and stained cells were imaged at 20x with widefield imaging modality on PICO ImageXpress (Molecular Devices), with 9 fields, using 2 z-sections combined with best focus for small objects function to ensure cells in focus across the entire field to accommodate deviations in the flatness of imaging plate surfaces. Acquired images were analyzed using IN Carta (Molecular Devices) analysis software. Nuclei were identified based on the Hoechst signal using an off-the-shelf SINAP model, and cell/cytoplasmic boundaries defined using the weak cellular staining with Hoechst with only one nucleus. TDP-43-GFP puncta and G3BP+ stress granules were defined using a robust puncta algorithm in the corresponding channels and assigned to the cytoplasm of individual cells for enumeration. For quantification, over 300 cells were analyzed per well to generate an average value. To normalize to % max for TDP-43-GFP puncta and G3BP1+ puncta, the average of the arsenite control (water-treated) wells was subtracted from the value of each well (defining 0%), followed by division with the average of the arsenite-treated DMSO-vehicle treated wells as the maximum response set to 100%. To normalize for nuclear TDP-43, 0% was defined by arsenite-treated and 100% by arsenite control wells.

#### Mis-splicing of TDP-43 targets in iPSC-derived motor neurons

Differentiated iPSC-derived motor neurons from a healthy individual were purchased from FUJIFILM and cultured under manufacturer directions. Briefly, 96-well imaging plates were pre-coated first with poly-D-lysine (Gibco) overnight, washed without drying, then double-coated with Geltrex (Ready-use, Gibco) for at least 1 hour before plating cells at 27-32,000 cells/well. Media was changed every 2-3 days, with 75% media changes with DAPT (Sigma-Aldrich) for the first week, and 50% media changes after that. After 2-3 weeks in culture from the date of plating, cells were treated with DMSO or ACE-339 for 24 hours. Sodium arsenite (or water control) was spiked into the culture media to 250µM for 120 minutes and cell lysates were harvested with TRI Reagent (Zymo). RNA was extracted and purified with Direct-zol RNA kits (Zymo) including on-column gDNA digestion. cDNA libraries were generated with SuperScript IV VILO Master Mix (Invitrogen) on a thermocycler. qPCR reactions were made using PowerTrack SYBR Green Master Mix (Thermo) in a 384-well format. Custom DNA primers (IDT) were chosen based on published literature (Table S2). qPCR was measured and analyzed using a QuantStudio 6 Pro Real-Time PCR System and Design & Analysis Software 2.7.0 software (Thermo). The relative quantification analysis setting was used with normalization to GAPDH and reference sample specified as the DMSO no arsenite condition to determine the fold change.

### iTDP-43 mouse model study

#### Mice

iTDP-43^A315T^ female and male mice were group housed on a 12h light/dark cycle with *ad libitum* access to food and water. One group of mice (DOX) was obtained from breeding pairs fed food containing doxycycline 200mg/kg (SF11-059 Specialty Feeds, Australia), and maintained on this food, turning off expression of the transgene for the duration of the experiment. No other mice were fed doxycycline food at any time.

#### Drug Treatment

Drug treatments were conducted in a double-blinded manner. ACE-339 and vehicle were formulated weekly for oral gavage delivery as follows: Vehicle consisted of Kolliphor EL (Thermo-Fisher Scientific, USA) mixed with water for injection (Baxter, USA) at a ratio of 1:4 v/v. Two concentrations of ACE-339 were dissolved in the vehicle. Briefly, the compound was dissolved by warming the solution to 40-45°C and sonicating, pH was adjusted to 3.5-4 using 1N HCl. The final formulation was aliquoted and stored at -20°C. Mice (except for the DOX group) were randomly assigned to groups: vehicle, ACE-339 75mg/kg or 150mg/kg and were dosed via oral gavage (p.o.) daily from 4 weeks of age for 56 days by a blinded experimenter not involved in preparation of the drugs. Mice were weighed twice weekly for the duration of the dosing period and were subjected to behavioral testing 4 and 8 weeks after commencing dosing.

#### Tissue Collection

For tissue collection, mice were anaesthetized 2h after the last dose at day 56 and blood collected by cardiac puncture, followed by transcardial perfusion with phosphate buffered saline. Brains were collected and halved, one hemisphere was fixed in 4% paraformaldehyde (PFA) overnight before histological processing, and the other hemisphere sub-dissected and snap-frozen in liquid nitrogen for protein analysis. All animal experiments were approved by the Macquarie University Animal Ethics Committee. All behavioral testing was conducted blinded to experimental group. Testing was performed by an experimenter not involved in preparation or administration of drugs.

#### Rotarod

Motor performance was determined at 8-weeks post initiation of drug treatment using a Rota-Rod (Ugo Basile) in acceleration mode (5–60 rpm) over 120s as previously described^65^. The longest time each mouse remained on the turning wheel out of five attempts was recorded.

#### Pole Test

To test strength and coordination at 4- and 8-weeks post initiation of drug treatment, mice were placed at the top of a vertical pole (47.5 cm length of dowel, 0.8 cm diameter) facing upwards and allowed to turn and descend as previously described^66^. The best time out of two turns taken to turn and descend was chosen. Mice unable to complete the test (slipped or fell) were given the maximum score of 120s.

#### Elevated Plus Maze

At 8-weeks post initiation of drug treatment, mice were acclimatized in the room for 1hr prior to being placed on the Elevated Plus maze (Ugo Basile), consisting of two open and two closed arms (each 35cm × 5.5cm), as well as a central platform (5.5cm × 5.5cm), elevated 60 cm above the ground, as previously described^67^. Mice were recorded for 5 min and videos analyzed using AnyMaze software.

#### Open Field

Activity and exploratory behavior were assessed at 8-weeks post initiation of drug treatment using 40cm square open field arena as previously described^68^. Videos were recorded and analyzed using AnyMaze software, with an inner zone of 17.5cm square in the center of the arena.

#### Pharmacokinetics

ACE-339 levels were measured in snap-frozen hemibrains using LC-MS/MS (Meadowhawk Biolabs). Brain samples were processed using an Omni beadruptor homogenizer following an addition of 4 eq (w/v) of 80:20 Water:Acetonitrile. Samples were analyzed against a matrix matched curve.

#### Sequential extraction

Sequential extraction with RIPA followed by urea buffers of cell and brain extracts was done as described^48^. 50–100mg of snap-frozen cortical tissue was solubilized in modified RIPA buffer (50mM Tris–HCl pH 8.0, 150mM NaCl, 1 % NP-40 substitute, 0.5% sodium-deoxycholate, 0.1% sodium dodecyl sulfate, 5mM EDTA, 30mM sodium fluoride, 60mM β-glycerophosphate, 20mM sodium pyrophosphate, 1mM sodium orthovanadate and 1x complete protease inhibitor (Roche)) using a motorized tissue douncer. The homogenate was sonicated at 20% amplitude for 30 short pulses, incubated at 4°C for 30 mins while rotating, then centrifuged at high-speed 50,000×g, at 4°C for 1 hour. The resulting supernatant constituted the RIPA-soluble fraction. To clear any remaining soluble proteins, pellets were washed twice by resuspending in RIPA buffer, followed by homogenization and high-speed centrifugation as above. The final pellet was resuspended in urea buffer (7M urea, 2M thiourea, 4% CHAPS in 30mM Tris, pH 8.5), sonicated at 20% amplitude for 30 pulses and centrifuged at 5,000xg for 10 mins to remove any cell debris. The resulting supernatant constituted the RIPA-insoluble/UREA soluble fraction. Sequential extraction of nuclear and cytoplasmic protein fractions was performed similarly to as previously described^69^. 50-100 mg cortical tissue was homogenized in CLB buffer (10mM HEPES, 10mM NaCl, 1mM KH_2_PO_4_, 5mM NaHCO_3_, 5mM EDTA, 1mM CaCl_2_, 0.5mM MgCl_2_ and 1x complete protease inhibitor (Roche)) using a motorized tissue douncer, then 0.1 volume 2.5M sucrose added for isotonic restoration (crude cell homogenate). Samples were then centrifuged at 6,300xg for 10 mins and pellets resuspended in TSE buffer (10mM Tris pH 7.2, 300mM sucrose, 1mM EDTA, 0.1% IGEPAL-CA 630 v/v), and centrifuged at 4,000xg for 5 mins, after which the supernatant was discarded, and the pellet washed in TSE buffer 3 times until supernatant was clear. The resulting pellet was resuspended in RIPA buffer, centrifuged at 14,000xg for 10 mins, and supernatant collected as RIPA soluble nuclear fraction. The pellet was resuspended in urea buffer and centrifuged at 14,000xg for 10 minutes, and collected as RIPA insoluble nuclear fraction. Supernatant from the first centrifugation step was sedimented at 6,300xg for 10 mins and the resulting supernatant collected (cytoplasmic fraction). Protein concentrations were determined using the Bradford assay (BioRad). Samples were aliquoted, snap frozen and stored at -80°C until further use.

#### Western blotting

Protein extracts were analyzed by western blotting, as described^70^. Primary antibodies used for immunoblotting were human TDP-43 (ProteinTech; 60019-2-IG; 1:2,000), pan-Ubiquitin (Dako; Z0458; 1/1000), pTDP-43 S409/410 (Cosmo Bio.; CAC-TIP-PTD-P02; 1/2000) and GAPDH (Merck-Millipore; MAB374; 1:5,000).

#### Immunohistochemistry

Fixed brains were processed using an automated system (Excelsior, Epredia USA), embedded in paraffin and sectioned at the level of the mid-hippocampus at 3μm thickness using a rotary microtome (Thermo-Microm USA). All staining was performed according to previously published protocols^71^. Primary antibodies used are as follows: human TDP-43 (ProteinTech; 60019-2-IG; 1:400), Iba1 (Wako; 019-19741; 1:500), Ctip-2 (Abcam; ab18465; 1:300), and GFAP (Sigma; G9269; 1:2,000). Secondary antibodies used were AlexaFluor-conjugated 488 and 555 (Thermo-Fisher Scientific; 1:250). Sections were scanned with an Axio Scan Z1 automated slide scanner (Zeiss). Quantification was done by an investigator blinded to the experimental groups. Cell counts were performed in a semi-automated manner using Zeiss Zen 2.6 Blue software and ImageJ, with macros and set parameters for each antibody.

### rNLS8 mouse model study

This study was conducted at PsychoGenics Inc following institutional IACUC guidelines.

#### Mice

Female and male rNLS8 mice were bred, group housed with 3-4 mice/cage in opti-MICE ventilated cages (Animal Care Systems, CO) and maintained at 12/12 light/dark cycles at 20-23°C and 50% relative humidity. Chow and water were provided *ad libitum* for the duration of the study. Mice were kept on doxycycline (DOX) diet until 5 weeks of age at which point mice were switched to regular diet. Testing was conducted during the light cycle of the mice. Mice in the same cage received the same treatment. All animals were examined and weighed prior to initiation of the study to assure adequate health and suitability. Body weights were measured twice per week throughout the study and mice were assessed for survival daily.

#### Drug treatment

ACE-339 was formulated as a suspension in 0.5% methylcellulose + 0.5% Tween80 in water at stored at 4 degrees C throughout the study. ACE-339 was administered orally once daily starting at 5 weeks of age immediately after DOX removal. Dosing continued till the end of the survival study.

#### Tissue collection and PK

For Nf-L analysis, plasma was collected from submandibular bleeds pre dosing and at 3 and 4 weeks post dosing. Blood was collected in tubes containing K2EDTA and was centrifuged for 10 min at 10,000-12,000 RPM, plasma collected and stored at -80° C. A satellite group of mice (n=4-5 per group) was used for tissue analysis after 4 weeks of dosing. Mice were anesthetized with isoflurane, perfused with PBS, plasma and brain tissue collected and frozen at -80C. ACE-339 drug levels were measured in plasma and cerebellum from the satellite group using LC-MS/MS (Quintara Discovery/Frontage Labs) as described above.

#### Nf-L measurement

Plasma was assessed for Nf-L using Quanterix SIMOA assay (PsychoGenics Inc. CAT#104073) using manufacturer’s protocol. Nf-L concentration was extrapolated from the constructed standard curve and corrected for dilution factor.

#### Homogenization and Sarkosyl Fractionation of TDP-43

This protocol was adapted^72^ with slight modifications. Frozen brain tissues were homogenized at a 1:5 weight/volume (w/v) ratio in high-salt (HS) buffer (10 mM Tris-HCl, pH 7.5; 0.8 M NaCl; 1 mM EGTA; 1 mM DTT) supplemented with 1X Halt™ Protease and Phosphatase Inhibitor Cocktail (Thermo Fisher Scientific, Cat# 78440). Homogenization was performed using a handheld cordless pestle motor homogenizer, followed by incubation on ice for 20–30 minutes. Sarkosyl (10%, Bioworld, Cat# 40120977-1) was added to the homogenate to a final concentration of 1% (v/v), and samples were incubated at 37°C for 30 minutes on a rotator. Lysates were centrifuged at 12,000 × g for 10 minutes at 16– 18°C, and a portion of the supernatant was collected as the total protein fraction. The remaining supernatant was ultracentrifuged at 100,000 × g for 30 minutes at 16–18°C to isolate the sarkosyl-insoluble (aggregated) protein pellet. This pellet was resuspended in 8 M urea/50 mM Tris-HCl, pH 8.0 with inhibitors and stored at −80°C until analysis.

#### Simple Western Analysis

Quantification of total and aggregated (sarkosyl-insoluble) hTDP-43 was performed at a certified external CRO (SBH Biosciences) under Bio-Techne’s quality standards using the Simple Western system (ProteinSimple). Protein concentrations were determined using a NanoDrop spectrophotometer, and equal amounts of protein were loaded into each capillary to ensure consistency across samples. Samples were denatured and prepared according to the manufacturer’s instructions for capillary-based immunodetection. Detection of human TDP-43 was performed using a human-specific anti-TDP-43 antibody (ProteinTech, Cat# 60019-2-Ig) at a dilution of 1:50. Aggregated hTDP-43 was defined as the signal obtained from the sarkosyl-insoluble (urea-solubilized) fraction, while total TDP-43 included signals from both the soluble and insoluble fractions. TDP-43 concentrations were calculated based on the sample signal and interpolation against a recombinant full-length human TDP-43 protein standard curve. Data were analyzed using Compass software (ProteinSimple).

### Molecular Dynamics simulations

#### Identifying the target hexameric structure of TDP-43

REMD simulations of the TDP-43 (307-319) peptide (referred to as ‘pre-formed TDP-43 hexamer’) were performed by randomly inserting six WT polypeptide structures in aqueous solution (100 mM NaCl buffer). Two separate REMD simulations were performed with different initial structures for the peptide: A) an extended conformation, B) a hairpin-like structure as obtained in earlier studies.^73^ Both the simulations were performed for 1100 ns, resulting in a combined 2200 ns of REMD simulation. The last 300 ns trajectories at 300 K temperature from each of these two simulations (600 ns in total) were used for analyzing data. Using the Daura algorithm, similar peptide structures were clustered based on their backbone root-mean-square deviation (RMSD), resulting in a total of 225 clusters^74^. The average hexameric structure from the most probable cluster was used as the target structure for the small molecule studies simulations.

#### Unbiased MD simulations

MD simulations were conducted by placing the ‘pre-formed WT’ hexamer in a dodecahedron box of 13 nm linear dimension. ACE-339 molecular geometries were subsequently inserted at random positions. The simulation box was solvated with 49,000 water molecules. 88 sodium ions and 94 chloride ions were added, to maintain electroneutrality at 100 mM salt concentration. 600 ns long trajectories were generated using a leap-frog integrator with 2 fs time steps^75^. The Charmm36m force fields for the peptides and the drug molecules were used in conjunction with the Charmm-modified TIP3P water model ^76,77^. NpT simulations were carried out at 300 K and 1 bar using the Nose-Hoover thermostat (1 ps relaxation time)^78,79^ and the Parrinello-Rahman barostat (3 ps relaxation time)^80^. Each simulation was preceded by 5 ns long equilibration simulations using the Berendsen thermostat (3 ps relaxation time) and barostat (3 ps relaxation time) at 300 K and 1 bar^81^. The bonds of the peptide and the drug molecules were constrained by the LINCS algorithm^82^. The bonded geometry of the water molecules was kept rigid using the SETTLE algorithm^83^. The nonbonded Lennard-Jones interactions were cut-off at 1.2 nm with long-range dispersion correction for pressure and energy. The particle mesh Ewald (PME) method with a real-space cut-off of 1.2 nm and a grid-spacing of 0.12 nm was used for calculating the electrostatic interactions^84^. The Gromacs (version 2021.2) package was used for the simulations^85^.

#### REMD simulations

Simulations were started by inserting six TDP-43 polypeptides at random positions in a dodecahedron box of 8 nm of linear dimension. Six ACE-339 molecules were inserted at random positions. The simulation box was solvated with 12,000 water molecules. 22 sodium ions and 28 chloride ions were added, to maintain electroneutrality at 100 mM salt concentration. 5 ns long NpT simulations were carried out using the Berendsen thermostat (0.5 ps relaxation time) and barostat (3ps relaxation time) at 300 K and 1 bar^81^. The compressibility value was set at 4.5*×*10*^−^*^5^ bar*^−^*^1^. Using the converged box-size corresponding to 1 bar pressure at 300 K, replica-exchange molecular dynamics (REMD) simulations were performed using 64 replicas at NVT ensemble^86–88^. The temperatures of the replicas were set at 292.7, 294.5, 296.3, 298.1, 300.0, 301.9, 303.8, 305.7, 307.7, 309.7, 311.7, 313.7, 315.8, 317.9, 320.0, 322.1, 324.3, 326.5, 328.7, 330.9, 333.2, 335.5, 337.8, 340.1, 342.5, 344.9, 347.3, 349.7, 352.2, 354.7, 357.2, 359.7, 362.3, 364.9, 367.5, 370.1, 372.8, 375.5, 378.2, 380.9, 383.7, 386.5, 389.3, 392.1, 395.0, 397.9, 400.8, 403.7, 406.7, 409.7, 412.7, 415.8, 418.9, 422.0, 425.2, 428.4, 431.6, 434.9, 438.2, 441.6, 445.0, 448.5, 452.0, and 455.6 K. The replicas were exchanged every 3 ps. Uniform exchange probabilities across all the temperatures with average probability of 20% to 30% were observed. The Gromacs (version 2021.2) package was used for the simulations^85^. The Nose-Hoover thermostat with 3 ps relaxation time was used for the REMD simulations^78,79^. The bonds of the peptide and the ligand molecules were constrained by the LINCS algorithm^82^. The bonded geometry of the water molecules was kept rigid using the SETTLE algorithm^83^. The nonbonded Lennard-Jones interactions were cut-off at 1.2 nm with long-range dispersion correction for pressure and energy. The particle mesh Ewald (PME) method with a real-space cut-off of 1.2 nm and a grid-spacing of 0.12 nm was used for calculating the electrostatic interactions^84^. A leap-frog algorithm and 2 fs time steps were used for integrating Newton’s equations of motion^75^. 1100 ns long trajectories were generated for each replica.

#### Data analyses

The peptide structures were clustered using the Daura algorithm where a 1.0 nm cutoff was used for the RMSD of the backbone N *−* C*_α_ −* C atoms. The ligand contacts with the peptides were determined using a 0.4 nm distance cutoff between the centers-of-mass of the ligand molecules (or the molecular groups) and centers-of-mass of the peptide atoms. For the calculations of the relative orientations of the peptides, peptides were assumed to be in contact if there were at least 100 interpeptide atomic contacts. The cutoff for the corresponding interpeptide contacts was determined from the REMD simulation of a WT dimer and the average number of the interpeptide contacts for the five most probable dimer configurations was used.

## Acknowledgements

We thank Kevin Burk for data management, Dr. Cody Aldaz for further developing the ML models and Dr. Andrew Carr for helpful discussions and the Acelot team for reviewing the manuscript. We thank our scientific advisors: Prof. Sami Barmada, Prof. Aaron Gitler and Prof. Gene Yeo, Dr. Marcie Glicksman, Dr. Mike Kavanaugh, Dr. Andrew J Perlman for helpful discussions and reviewing the manuscript. We thank Prof. Carol Hall and Xin Dong for additional *in silico* work not presented here. We thank the team at the Macquarie Animal Research Services, Macquarie University for animal husbandry. We thank the University of Edinburgh (Prof. Colin Smith) and Georgetown University (Dr. Brent Harris) for providing human postmortem tissue samples used for TDP-43 quantitation. We gratefully acknowledge that the Postmortem Tissue Cores at the University of Edinburgh and Georgetown University received funding from Target ALS to facilitate ALS bio-banking and ALS research. We thank the Institute for Brain Aging and Dementia at UC Irvine for providing human postmortem tissue samples used to isolate TDP-43 oligomers. We express our profound gratitude to the donors and their families whose generous contributions made this research possible.

## Author Contributions

VM, LW, and MK designed and oversaw the studies and wrote the manuscript. MK, SA, DK, VS, ES conducted recombinant TDP-43 and cellular assays. MAH, NM, AL, RK conducted the TDP-43 human brain derived oligomer experiments. AVH, MS, YDK conducted the iTDP43 mouse study. PG, JES, AS designed and conducted molecular dynamics simulation studies. All authors reviewed the manuscript.

