## Supplement for "In silico designed small molecule targets misfolded TDP-43 to rescue function and ALS pathology"

### Supplementary information

#### Methods

##### Full-length TDP-43 ThT

Recombinant full-length TDP-43 with His-tag at the N-terminus used in ThT assay was custom-made at Genscript. This material was used for the MST spectral shift experiments and dot-blot experiments as well. Purified full-length TDP-43 protein was eluted in urea and concentrated using a 10 kDa molecular weight cut-off filter. The protein was then refolded in a native buffer composed of 1X PBS, resulting in a final urea concentration of approximately 300 mM. For Thioflavin-T (ThT) fluorescence measurements, four solutions were prepared: the ACE-339 treatment group contained 5 μM native TDP-43, 150 μM ACE-339 and 30 μM ThT (VWR 103011-384). The vehicle control had 5% DMSO instead of compound, and the background control were the same except lacking ThT. The experiment was conducted in a 96-well plate with three replicates per sample. Fluorescence intensity measurements were recorded in kinetics mode every 15 minutes over a 72-hour period at 37C using a Synergy Neo2 plate reader, with excitation at 440 nm and emission at 490 nm. The intensity values of the averages of the 3 background control wells lacking protein were subtracted from the wells that include protein to remove background fluorescence signal from the plate, buffer, and TDP-43 protein. Data were exported as CSV files for analysis and plotted using GraphPad Prism software.

##### TDP-43 Low complexity domain (LCD) ThT

Recombinant TDP-43 LCD fragment with His-tag at the N-terminus was custom-made at Genscript. Purified TDP-43 LCD was eluted in urea and subsequently refolded in 1X PBS buffer (pH 7.4), achieving a final urea concentration of approximately 200 mM. For Thioflavin-T (ThT) fluorescence assays, three distinct solutions were prepared: the ACE-339 treatment group comprised 150 μM ACE-339 and 30 μM ThT along with 5 μM TDP-43 LCD. Both the vehicle control and the blank contained 5% DMSO and 30 μM ThT, with the blank solution using refolding buffer without protein. The experiments were performed in a 96-well plate format, with three replicates for each condition. Fluorescence intensity readings were taken every 10 minutes over a 24-hour duration using a Synergy Neo2 plate reader, with excitation set at 440 nm and emission detected at 490 nm. The resulting data were exported as CSV files for further analysis and plotting using GraphPad Prism software.

**Dynamic Light Scattering (DLS)**

Recombinant full-length TDP-43 (Genscript, supplied in 8M urea, 50mM Tris buffer) was introduced into native buffer (1X PBS, 5mM DTT) by fast dilution to a final concentration of 7μM. DLS measurements were performed immediately after dilution (0h) and 1.5 hours later to monitor aggregate formation. Data were collected using a Wyatt DynaPro instrument with 36 acquisitions of 10 seconds each at room temperature. Data were analyzed using the Dynamics software.

**Tubulin polymerization**

ACE-339 was tested in a tubulin polymerization assay to assess whether it might non-specifically disrupt the beta-sheet structure or affect functional polymerization of proteins, such as the formation of tubulin microtubules (Tubulin Polymerization Assay Kit BK011P, Cytoskeleton, Denver, CO). This kit contains a dye that increases fluorescence when incorporated into microtubules. The protocol provided with the kit was followed with slight modifications. Briefly, all components were reconstituted or thawed as per the manufacturer's instructions, and reagents were maintained on ice until the assay was initiated. A reaction mixture containing 2 mg/mL tubulin and 1 mM GTP was prepared for the polymerization reaction. The effects of ACE-339 on tubulin polymerization were assessed at eight concentrations ranging from 200 μM to 100 nM to evaluate a dose-response relationship. Vinblastine was used as a positive control at the same concentrations to inhibit tubulin polymerization. For the no-tubulin control, a supplemented buffer A was used in place of tubulin, and 2% DMSO served as the vehicle control. The assay was performed in fluorescence mode using a Neo2 Synergy plate reader (excitation wavelength: 360

nm, emission wavelength: 420 nm) in 96-well half-area plates (Corning, New York, NY). Data were exported in CSV format and analyzed using GraphPad Prism software.

**Microscale Thermophoresis**

Microscale thermophoresis (MST) experiments were conducted using a Monolith X (red-continuous) instrument (Nano Temper Technologies) to assess the binding affinity between full-length TDP-43 and ACE-339. Full-length TDP-43 was labeled with the Monolith protein labeling kit RED-MALEIMIDE (Nano Temper Technologies) under denaturing conditions (50 mM Tris pH 7.5, 8 M urea, 0.005% Tween-20, 5% DMSO) at a 1:3 protein ratio. Following labeling, free dye was removed using a PD Minitrap™ G-25 column (GE Healthcare). The eluted labeled protein was refolded by diluting it to 10 nM in a native buffer (1× PBS pH 7.4, 0.5 mM DTT, 0.005% Tween-20, 1% DMSO) immediately prior to the experiment. Note this is 500-fold more dilute than what was needed for TDP-43 to aggregate over time. Samples were loaded into Premium Coated Capillaries (NanoTemper Technologies) by capillary action. Measurements were performed at 25°C with 100% LED power and 40% laser power. ACE-339 was titrated from 100 μM to 3 nM in 16 serial 1:1 dilution steps with 10 nM of labeled TDP-43. A TDP-43 rabbit polyclonal antibody was used as a positive control, also titrated with 10 nM labeled TDP-43 in 16 serial 1:1 dilution steps from 1 μM to 30 pM. The normalized fluorescence was plotted against ligand concentrations. Data were analyzed and fitted using a non-linear regression model in GraphPad Prism to determine Kd values.

**SH-SY5Y cellular assay**

SH-SY5Y human neuroblastoma cells were obtained from ATCC (CRL-2266) and maintained in 1:1 EMEM/ F12 media supplemented with 10% FBS. Cells were seeded at 2500 cells per $cm^2$ in 12 well plate and were allowed to undergo differentiation into neuron-like cells by treating with 10 μM all-trans retinoic acid (Sigma-Aldrich R2625-50MG) for 6 days in EMEM/F12 media supplemented with 10% FBS, followed by 4 days of treatment with 50 ng/mL of BDNF (Peprotech

450-02B) in EMEM/F12 media. On day 10, half of the media was replaced by 50ng/ml of BDNF containing EMEM/F12 media and three different doses of ACE-339 were added such that the final concentrations were 10uM, 30uM and 100uM. After 20-22 hours of incubation with ACE-339, 250uM of sodium arsenite solution was added for 2 hours. Cells were harvested after 2 hours with Trizol reagent for RNA extraction. Total RNA was extracted using Direct-zol RNA Miniprep Kit (Zymo Research R2070). RNA was quantified by Nanodrop (ThermoFisher Scientific), and 300 ng RNA was used for cDNA synthesis with SuperScript IV VILO Master Mix (ThermoFisher Scientific 11756050). Quantitative PCR was run with 5 ng cDNA input in a 20 µl reaction using PowerTrack SYBR Green Master Mix (ThermoFisher Scientific A46109) with readout on a QuantStudio 6 Flex (ThermoFisher Scientific) using standard cycling parameters (95 °C for 2 min, 40 cycles of 95 °C for 15s and 60 °C for 60 s), followed by standard dissociation (95 °C for 15 s at 1.6 °C s−1, 60 °C for 60 s at 1.6 °C s−1, 95 °C for 15 s at 0.075 °C s−1). ΔΔCt and fold changes (Rq) with respect to untreated condition were calculated with GAPDH as the housekeeper gene; measured Ct values greater than 40 were set to 40 for visualizations in the QuantStudio Design and Analysis software (ThermoFisher Scientific). The Rq values were exported in the csv format and the statistical analysis was performed in GraphPad Prism. See Table S2 for primers.

**TDP-43 quantification via Protein Simple Western™ capillary electrophoresis**

TDP-43 protein quantification in human post-mortem brain tissues were measured using semi-quantitative capillary electrophoresis method.

*Tissue collection*: Frozen postmortem brain tissues (motor cortex) were obtained from 14 sporadic ALS (sALS) patients and 8 neurologically healthy controls. The demographic and clinical characteristics of the cohort are detailed in Table S1. All tissues were sourced from Target ALS biobanks at Georgetown University and University of Edinburgh and collected under respective institutional ethical approvals. Samples were stored at −80°C until further processing.

*Homogenization*: Frozen brain tissues were homogenized at a 1:5 weight/volume (w/v) ratio in high-salt (HS) buffer (10 mM Tris-HCl, pH 7.5; 0.8 M NaCl; 1 mM EGTA; 1 mM DTT) supplemented with 1X Halt™ Protease and Phosphatase Inhibitor Cocktail (Thermo Fisher Scientific). Homogenization was performed using a handheld cordless pestle motor homogenizer, followed by incubation on ice for 20–30 minutes. Sarkosyl (10%, Bioworld) was added to a final concentration of 1% (v/v), and samples were incubated at 37°C for 30 minutes on a rotator. Lysates were centrifuged at 12,000 × g for 10 minutes at 16–18°C, and the supernatant was collected as the total protein fraction.

*Simple Western Assay*: Quantification of total TDP-43 in human tissues was performed at a certified external CRO (SBH Biosciences) under Bio-Techne's quality standards. Protein concentrations were determined by NanoDrop, and equal amounts of protein were loaded into each capillary to ensure consistency across samples. Since equal protein amounts were loaded and since total protein normalization performed on all patient samples yielded normalization factor of 1.0 on average, no further normalization was performed beyond correcting for the dilution and homogenization factor. This approach ensured that absolute TDP-43 concentrations were calculated based on the sample signal and interpolation against a recombinant full-length human TDP-43 protein standard curve, avoiding the introduction of additional correction factors. Detection employed a TDP-43 (C-terminal) polyclonal antibody (Proteintech, Cat# 12892-1-AP). For pTDP-43 measurements, a phospho-specific antibody targeting TDP-43 phosphorylated at S409/410 (rabbit monoclonal, clone 26H10 from Mayo Clinic (Zhang andPetrucelli labs), sourced from Target ALS through DSHB) was used. Data analysis was conducted using Compass software, with TDP-43 concentrations were calculated by fitting signals to a four-parameter logistic (4PL) regression model used to fit the standard curve.

**TDP-43 quantification via ELISA**

TDP-43 protein concentrations in mouse brain tissues were measured using the SimpleStep ELISA® kit for TDP-43 (Abcam, Catalog no. ab282880), following the manufacturer's protocol with minor modifications. Briefly, brain tissue homogenates were prepared by dissolving PBS-rinsed tissue in a volume of chilled 1X cell extraction buffer, which was supplemented with protease and phosphatase inhibitor cocktails at a volume equal to five times the weight of the tissue. The mixture was then homogenized with a handheld homogenizer and a disposable pestle. Samples were incubated on ice for 20 minutes, followed by centrifugation at 18,000 × g for 20 minutes at 4°C. The resulting supernatants were transferred to clean Eppendorf tubes, aliquoted, and stored at -80°C until further analysis. Total protein concentrations were determined using the Pierce BCA Protein Assay Kit (ThermoFisher Scientific 23225). For the ELISA, 50 µL of diluted sample or standard and 50 µL of antibody cocktail were added to the wells of microplate strips provided with the kit and incubated for one hour on a plate shaker set at 400 rpm. The plate was then washed three times with wash buffer as specified in the protocol, followed by the addition of TMB development solution. The development of the blue color was monitored by measuring absorbance at 600 nm in kinetics mode using a Neo2 Synergy plate reader. Once the absorbance at 600 nm reached approximately 0.8, the stop solution was added, and the absorbance at 450 nm was recorded. TDP-43 concentrations in the samples were determined by interpolating the blank control-subtracted absorbance values against a standard curve in the software provided with the Neo2 Synergy plate reader. Statistical analysis was performed in GraphPad Prism.

**Wild type mouse pharmacokinetic study**

This study was conducted at Enamine/Bienta following organizational IACUC. CD-1 male mice (8-9 weeks old) were housed in standard cages at 22+/- 3C, 40-70% relative humidity in 12h light/dark cycle. Mice were given standard rodent diet and water *ad libitum*, and were fasted 6 hours before each blood sampling. ACE-339 was formulated as a suspension using Kolliphor EL – water for injections (20%:80%). Mice were dosed orally with vehicle or different doses of ACE-339 daily for 14 days. Blood was collected by submandibular bleeds and plasma prepared for subsequent analysis was done at 0, 5, 15, 30, 60, 120, 240, 480, and 1440 min after the first and 14th dose. Separate groups of mice were dosed similarly, euthanized and perfused for brain collection and subsequent analysis at 60, 480 and 1440 mins after the 1$^{st}$ and 14$^{th}$ doses. Plasma samples (20 µl) were mixed with 100 µl of IS(90) solution. After mixing by pipetting and centrifuging for 4 min at 6,000 rpm. Brain samples (weight 113 mg – 202 mg) were homogenized with 5 volumes of IS(80) solution (1 w + 5 v) using zirconium oxide beads (115 mg ± 5 mg) in The Bullet Blender® homogenizer for 30 seconds at speed 8. After this, the samples were centrifuged for 4 min at 14,000 rpm, and 1 µl of each plasma and brain supernatant was injected into LC-MS/MS system.

**RNA-seq and Pathway Analysis**

RNA extraction, library preparation, sequencing, and bioinformatics analysis were performed by Eremid. Briefly, RNA was extracted from mouse brains (4 replicates per condition) using the Qiagen RNeasy Plus Universal Mini Kit and quality-assessed using TapeStation (Agilent) and Qubit 3.0 (Thermo Fisher Scientific); all samples had RIN ≥ 8.2. Libraries were prepared using the Universal Plus Total RNA-Seq Kit (Tecan) with ERCC spike-in controls and sequenced on an Illumina NovaSeq6000 (100 bp paired-end reads), generating approximately 230 million total paired-end reads per library. Raw reads were demultiplexed, trimmed, and aligned to the GRCm39 mouse reference genome, followed by read quantification and differential isoform expression analysis. Differential isoform expression between vehicle-treated (n=4) and ACE-339 400mg/kg-treated (n=4) groups were visualized using volcano plots generated in R (version 4.5.0) with the EnhancedVolcano package, applying thresholds of adjusted p-value $< 0.05$ and $|\log 2 \text{ fold change}| > 1$. Pathway enrichment analysis was performed by Eremid, and pathways relevant to the rNLS8 model were visualized in GraphPad Prism ($p < 0.05$).

**Molecular Dynamics simulations**

*Unbiased MD simulations:* 600 ns long trajectories were generated using a leap-frog integrator with 2 fs time steps (Verlet Phys Rev 1967). The Charmm36m force fields for the peptides and the drug molecules were used in conjunction with the Charmm-modified TIP3P water model (Huang Nat Methods 2017, MacKerell Mol Phys 1984). NpT simulations were carried out at 300 K and 1 bar using the Nose-Hoover thermostat (1 ps relaxation time) (Nose Mol Phys 1984,

Hoover Phys Rev A 1985) and the Parrinello-Rahman barostat (3 ps relaxation time) (Parrinello J Appl Phys 1981). Each simulation was preceded by 5 ns long equilibration simulations using the Berendsen thermostat (3 ps relaxation time) and barostat (3 ps relaxation time) at 300 K and 1 bar (Berendsen J Chem Phys 1984). The bonds of the peptide and the drug molecules were constrained by the LINCS algorithm (Hess J Comput Chem 1997). The bonded geometry of the water molecules were kept rigid using the SETTLE algorithm (Miyamoto J Comput Chem 1992). The nonbonded Lennard-Jones interactions were cut-off at 1.2 nm with long-range dispersion correction for pressure and energy. The particle mesh Ewald (PME) method with a real-space cut-off of 1.2 nm and a grid-spacing of 0.12 nm was used for calculating the electrostatic interactions (Essmann J Chem Phys 1995).

Five ns long NpT simulations were carried out using the Berendsen thermostat (0.5 ps relaxation time) and barostat (3ps relaxation time) at 300 K and 1 bar (Berendsen J Chem Phys 1984). The compressibility value was set at $4.5\times10^{-5}$ $bar^{-1}$. Using the converged box-size corresponding to 1 bar pressure at 300 K, replica-exchange molecular dynamics (REMD) simulations were performed using 64 replicas at NVT ensemble (Hukushima & Nemoto JPhys Soc Jpn 1996, Sugita Chem Phys Lett 1999, Patriksson Phys Chem Chem Phys 2008). The temperatures of the replicas were set at 292.7, 294.5, 296.3, 298.1, 300.0, 301.9, 303.8, 305.7, 307.7, 309.7, 311.7, 313.7, 315.8, 317.9, 320.0, 322.1, 324.3, 326.5, 328.7, 330.9, 333.2, 335.5, 337.8, 340.1, 342.5, 344.9, 347.3, 349.7, 352.2, 354.7, 357.2, 359.7, 362.3, 364.9, 367.5, 370.1, 372.8, 375.5, 378.2, 380.9, 383.7, 386.5, 389.3, 392.1, 395.0, 397.9, 400.8, 403.7, 406.7, 409.7, 412.7, 415.8, 418.9, 422.0, 425.2, 428.4, 431.6, 434.9, 438.2, 441.6, 445.0, 448.5, 452.0, and 455.6 K. The replicas were exchanged every 3 ps. Uniform exchange probabilities across all the temperatures with average probability of 20% to 30% were observed.

The Nose-Hoover thermostat with 3 ps relaxation time was used for the REMD simulations (Nose Mol Phys 1984, Hoover Phys Rev A 1985). The bonds of the peptide and the ligand molecules were constrained by the LINCS algorithm (Hess J Comput Chem 1997). The bonded geometry of the water molecules were kept rigid using the SETTLE algorithm (Miyamoto J comput Chem

1992). The nonbonded Lennard-Jones interactions were cut-off at 1.2 nm with long-range dispersion correction for pressure and energy. The particle mesh Ewald (PME) method with a real-space cut-off of 1.2 nm and a grid-spacing of 0.12 nm was used for calculating the electrostatic interactions (Essmann J Chem Phys 1995). A leap-frog algorithm and 2 fs time steps were used for integrating Newton's equations of motion (Verlet Phys Rev 1967). 1100 ns long trajectories were generated for each replica.

**Figure S1. ACE-339 specifically targets aggregated TDP-43.** (A) ThT dye binding assay indicates that ACE-339 dose dependently inhibits de novo TDP-43 (307-319) aggregation. The molar $IC_{50}$ for ACE-339 is dependent on TDP-43 concentration used in the assay (10, 30, 100 μM), but target to drug molar ratio remains constant irrespective of target load. (B-C) ACE-339 inhibits aggregation of recombinant LCD fragment (B) and TDP-43 full length protein (C) at a 1:30 protein:drug in ThT kinetic assay. (D) A light absorbance scan from 300-700nm. Compared to the DMSO control (gray), ACE-339 does not absorb significant light at the excitation wavelength of (440nm; box indicates 430-450nm) or the emission wavelength (490nm; box indicates 480-500nm) of ThT, excluding primary and secondary inner filter effect artifacts of ThT interference as a false positive for reduced ThT signal. (E) Dynamic light scattering (DLS) size distributions of full-length TDP-43 following rapid dilution into native buffer. Immediately upon dilution, the protein forms a discrete 7.6 nm hydrodynamic radius monomeric population (left). After 1.5 hours of incubation, this monomer transitions completely into large >130 nm radius aggregates (right). These data indicate that freshly diluted recombinant full-length TDP-43 source is monomeric prior to incubation and subsequent aggregation. (F) Microscale Thermophoresis (MST) fluorescence spectral shift showing ACE-339 does not bind to natively folded full length TDP-43 protein (left), while a N-terminal specific TDP-43 antibody control shows dose dependent binding (right). Dissociation constants, Kd, are calculated. n.d.= not determined. (G) Tubulin polymerization assay showing ACE-339 (and vehicle treatment) does not inhibit polymerization of unrelated functional protein. A dye increases fluorescence in the presence of microtubules. Vinblastine was used as a positive inhibitor control.

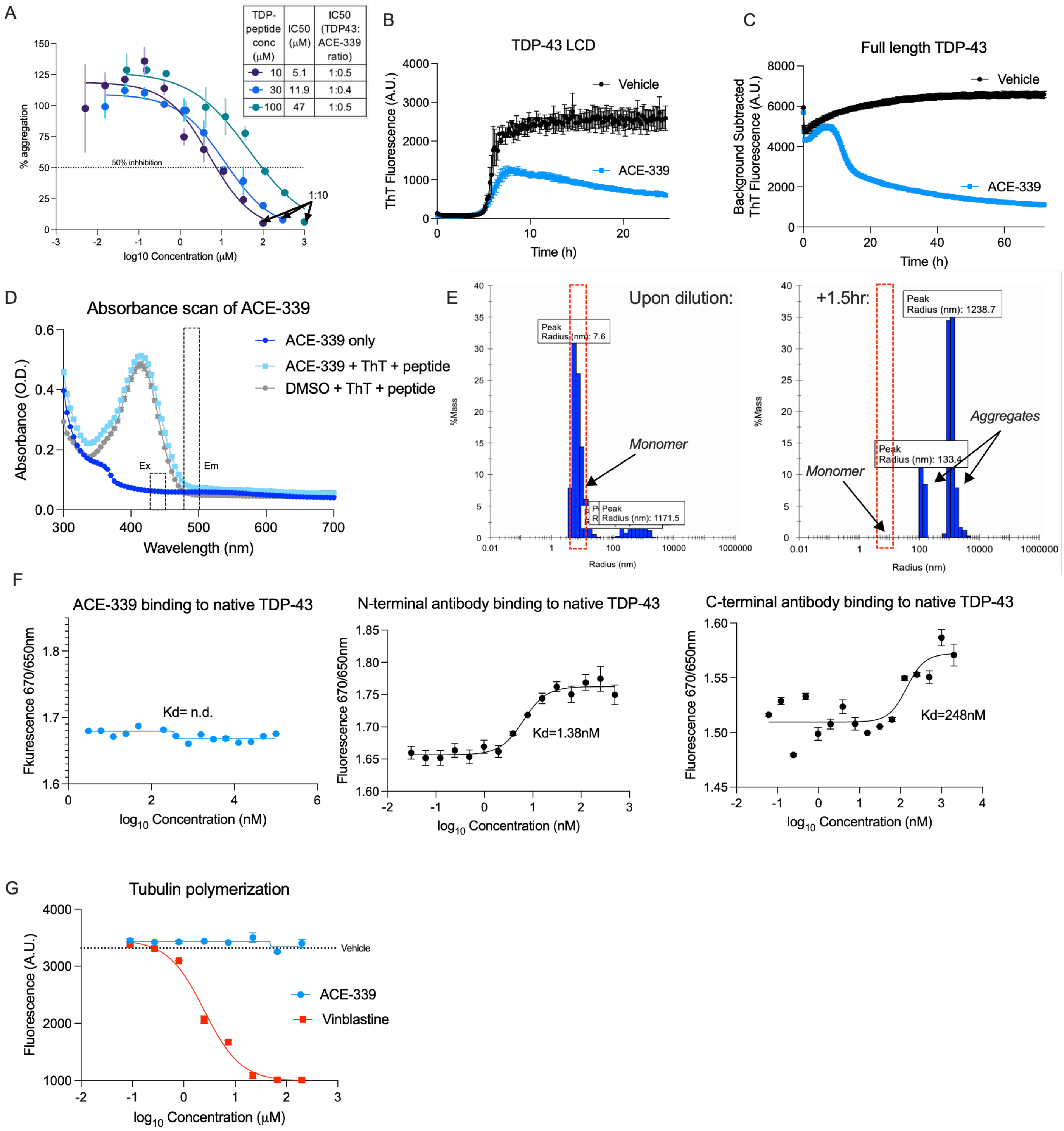
A
TDP-peptide conc (μM)
IC50 (μM)
IC50 (TDP43: ACE-339 ratio)
10 5.1 1:0.5
30 11.9 1:0.4
100 47 1:0.5
% aggregation
50% inhhibition
1:10
log10 Concentration (μM)
B
TDP-43 LCD
ThT Fluorescence (A.U.)
Vehicle
ACE-339
Time (h)
C
Full length TDP-43
Background Subtracted ThT Fluorescence (A.U.)
Vehicle
ACE-339
Time (h)
D
Absorbance scan of ACE-339
Absorbance (O.D.)
ACE-339 only
ACE-339 + ThT + peptide
DMSO + ThT + peptide
Ex
Em
Wavelength (nm)
E
Upon dilution:
Peak Radius (nm): 7.6
Monomer
Peak Radius (nm): 1171.5
%Mass
Radius (nm)
+1.5hr:
Peak Radius (nm): 1238.7
Aggregates
Peak Radius (nm): 133.4
Monomer
F
ACE-339 binding to native TDP-43
Fkurescence 670/650nm
Kd= n.d.
log10 Concentration (nM)
N-terminal antibody binding to native TDP-43
Fluorescence 670/650nm
Kd=1.38nM
log10 Concentration (nM)
C-terminal antibody binding to native TDP-43
Fluorescence 670/650nm
Kd=248nM
log10 Concentration (nM)
G
Tubulin polymerization
Fluorescence (A.U.)
Vehicle
ACE-339
Vinblastine
log10 Concentration (μM)

**Figure S2. ACE-339 has a novel mechanism of action**. 50μM TDP-43 (307-319) was coincubated with DMSO control or ACE-339 at 1:1, 1:2 and 1:5 molar ratios for 24h. A parallel experiment included ThT dye to confirm ACE-339 activity (left kinetic graph). Each reaction mix (non-ThT) was passed through 1kDa molecular weight cut-off filter to separate TDP-43 (307-319) bound and free drug. ACE-339 concentration and activity were measured in the "free drug" fraction (right graphs). 50-85% of the initial ACE-339 drug was detected in the free fraction (right, top). The drug free fraction, but not vehicle free fraction, inhibited the aggregation of newly added TDP-43 (307-319) monomers (right, bottom).

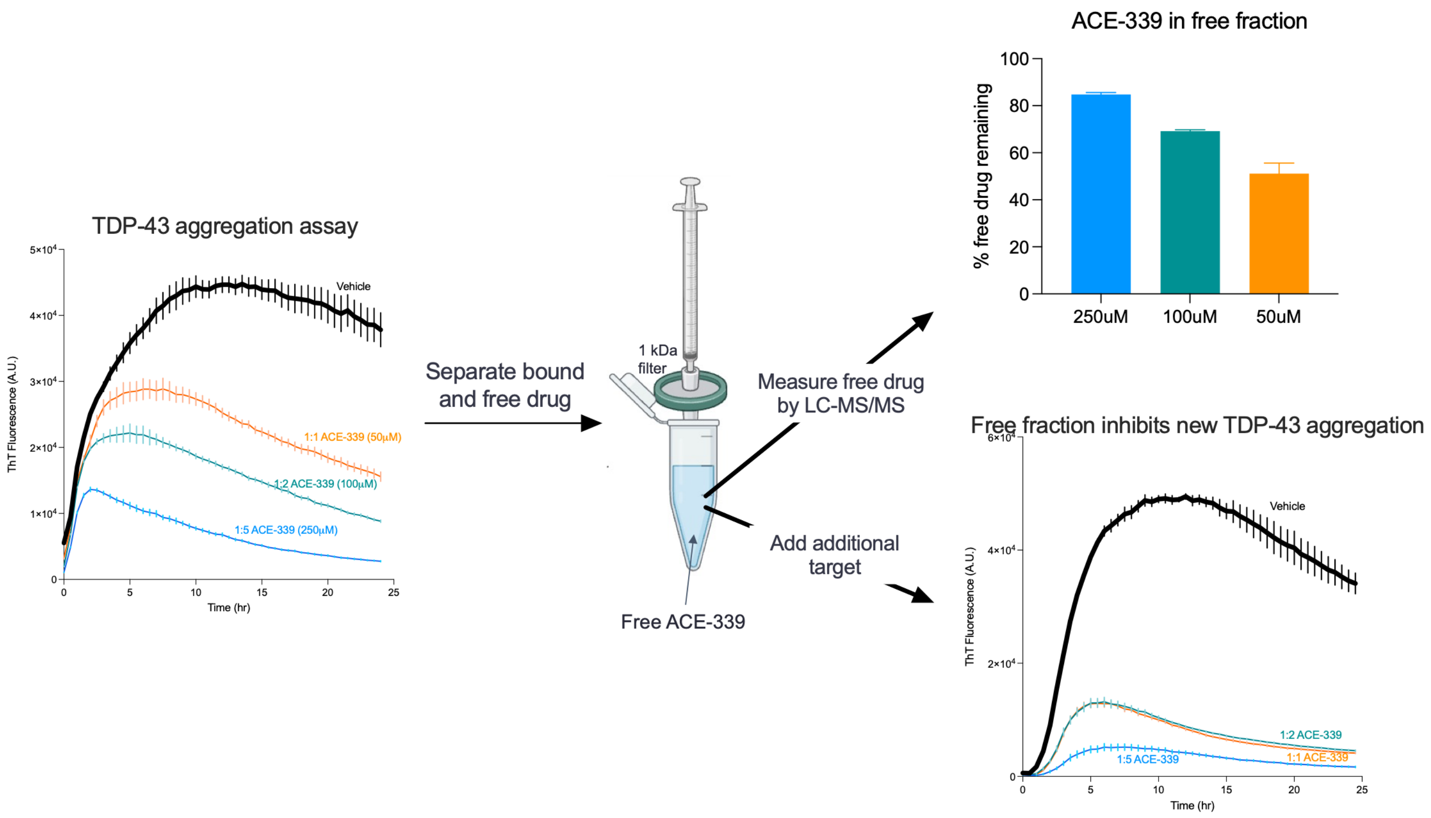

**Figure S3. ALS brain derived oligomer seeded recombinant TDP-43 conformations are similar to native ALS brain derived TDP-43 oligomers and can mature into larger fibrils.** (A) Oligomers that were purified from ALS patient brain tissue are positive for TDP-43 by immunoblot (IB). Samples were separated with SDS-PAGE and immunoblotted with an antibody specific to TDP-43. (B) Seeded recombinant TDP-43 oligomers show the same molecular high weight as the brain derived TDP-43 oligomers from which they were templated. To maintain native oligomeric complex sizes, samples were separated with Native PAGE and all protein detected by silver stain. While recombinant full length TDP-43 (right) displayed monomeric size, the original ALS patient brain derived TDP-43 oligomers (left) displayed a higher molecular weight. These were used to seed monomers, and the resultant oligomer seeded full length TDP-43 conformations (middle) displayed the same higher molecular weight as the original patient derived oligomers. Molecular weights in kilodaltons (kDa) are indicate to the left, and sizes consistent with hexamer and monomers of TDP-43 are on the right. For each condition, both 5 and 2.5 µg of sample was loaded into the gel. (C) ALS patient brain derived oligomer seeded full length TDP-43 conformations go on to form fibrils at longer time points. AFM images (top) and line traces (bottom) displaying length and height of ALS patient brain derived oligomer seeded full length TDP-43 conformations that formed fibrils spontaneously formed at 37 C and pH 7.4 by rotation at 10-15 rpm for 5 days, which is 3 days after oligomers form. All fibrils are of similar height and periodicity. The same fibril morphologies were observed in 4 images for each type of sample, and 2 representative regions are shown here, a 5µm x 5µm on the left and a 2µm x 2µm on the right. Colored arrows indicate which line trace is plotted below.

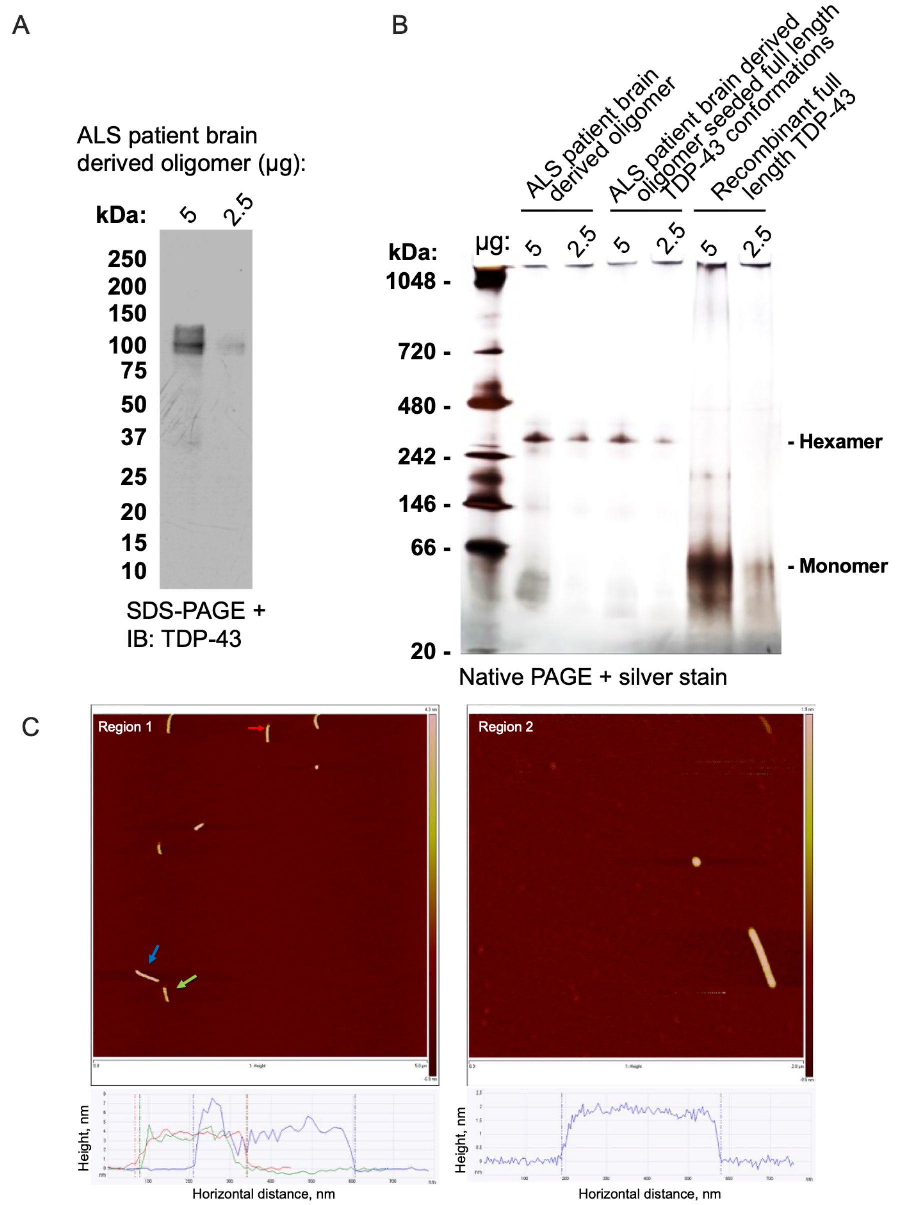
A
ALS patient brain derived oligomer (µg):
kDa:
5
2.5
250
200
150
100
75
50
37
25
20
15
10
SDS-PAGE + IB: TDP-43
B
ALS patient brain derived oligomer
ALS patient brain derived oligomer seeded full length TDP-43 conformations
Recombinant full length TDP-43
kDa:
µg:
5
2.5
5
2.5
5
2.5
1048 -
720 -
480 -
242 -
146 -
66 -
20 -
- Hexamer
- Monomer
Native PAGE + silver stain
C
Region 1
Region 2
Height, nm
Horizontal distance, nm
Height, nm
Horizontal distance, nm

**Figure S4. ACE-339 rescues TDP-43 oligomer induces synaptic defects.** Baseline transmission was calculated in slices treated with 100nM TDP43 oligomers (TDP-43O), TDP-43 oligomers pre-incubated with the anti-TDP43 oligomer drug ACE-339 at a 1:10 or 1:20 ratio, ACE-339 alone (0.5μM), or recombinant tau oligomers (TauO, positive control) by measuring fiber volley (FV) amplitude and field excitatory post-synaptic potential (fEPSP) slopes as the stimulus intensity is gradually increased from 100-1000μA over a 10 minutes period. (A) fEPSP slopes plotted against FV amplitudes reveal differential relationships between pre-synaptic engagement and postsynaptic output between treatment groups. TDP43O-treated groups reach a plateau in baseline responsiveness faster and with less presynaptic fiber-volley engagement than untreated, vehicle-treated (ACE-339), and TauO treated groups. Vehicle (n=7), ACE-339 Alone (n=3), TDP-43O+ACE-339 1:20 (n=7), TDP-43O+ACE-339 1:10 (n=4), TDP-43O (n=6), RTauO (n=5). IO-FV profiles were analyzed in GraphPad Prism 10 using the non-linear fit function Y=Y0+(Plateau-Y0)*(1-exp(-K*x)), where Y is the fEPSP slope, X is the fiber volley amplitude, and K is the rate constant. (B) Paired-Pulse ratio for all groups. (C-D) Long term potentiation profiles (D) and fEPSP slope quantification in the last 10 minutes (C) for all groups are shown. Mean±SEM are shown. P<0.05*, P<0.01**. One-way ANOVA followed by Kruskal-Wallis test.

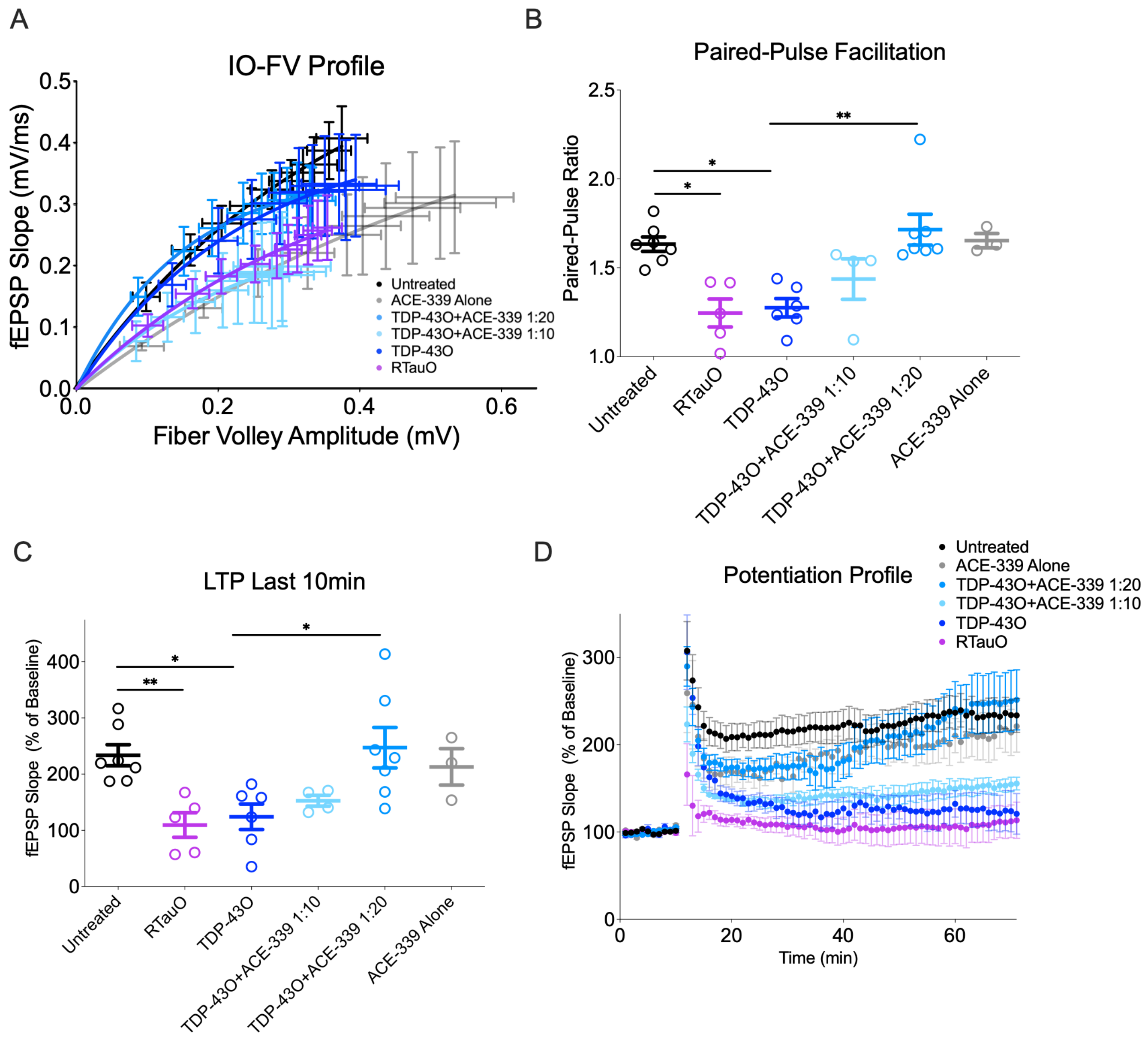

**Figure S5. ACE-339 rescues gain-of-function arsenite stress-free TDP-43-GFP cytoplasmic puncta in U2OS and loss-of-function cryptic exons inclusion in arsenite SH-SY5Y differentiated neuron model.** (A) U2OS cells with TDP-43-GFP overexpression treated with ACE-339 show reduced number of TDP-43-GFP cytoplasmic puncta in an arsenite stress-free model. (B) Differentiated SH-SY5Y cells treated with 250μM sodium arsenite with ACE-339 24h pretreatment. Data shows mRNA fold change in quantitative RT-PCR using specific primers for cryptic exons, representative of 2 independent runs with n=2-3 biological repeats. Mean±SEM are shown. $P<0.01$**, $P<0.0001$**** One-way ANOVA followed by Dunnett's (A) or Kruskal-Wallis (B) test.

A

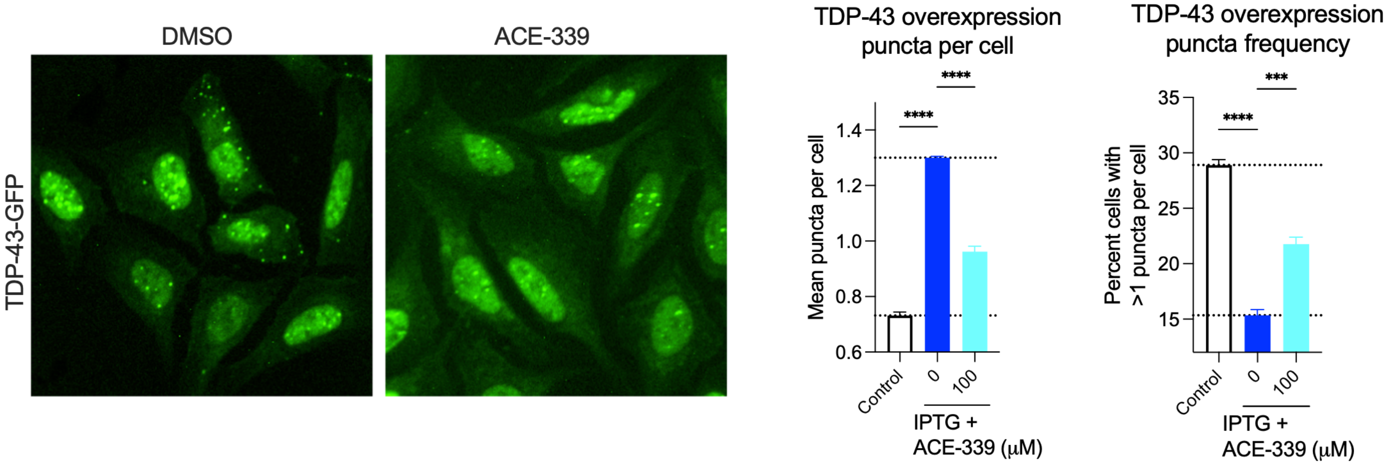


B

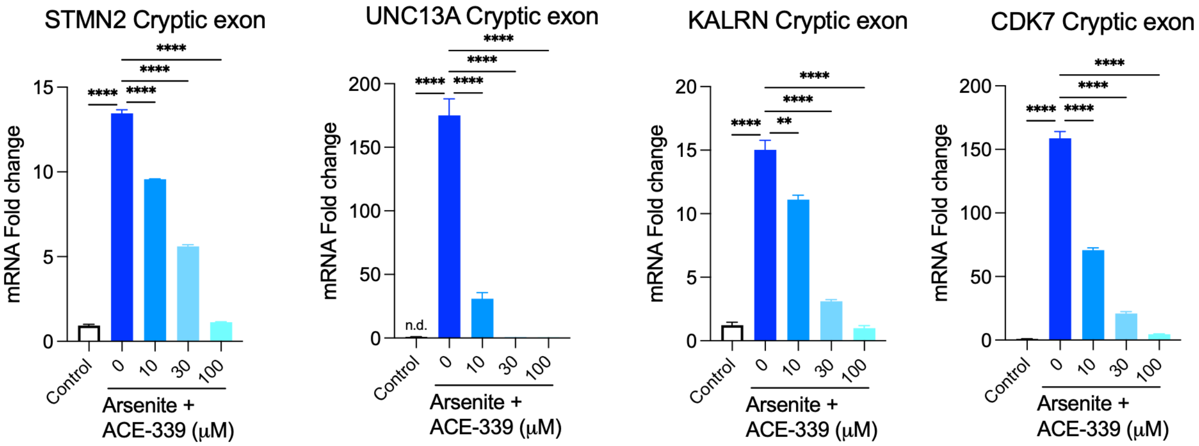

**Figure S6. ACE-339 pharmacokinetic (PK) properties in plasma and brain of wildtype mice.** ACE-339 was dosed orally once a day for 14 days in CD-1 male mice (n= 3/group). (A-B) Plasma and brain PK at indicated time points after dosing on day 1 and day 14 at indicated doses. Means are shown at each time point.

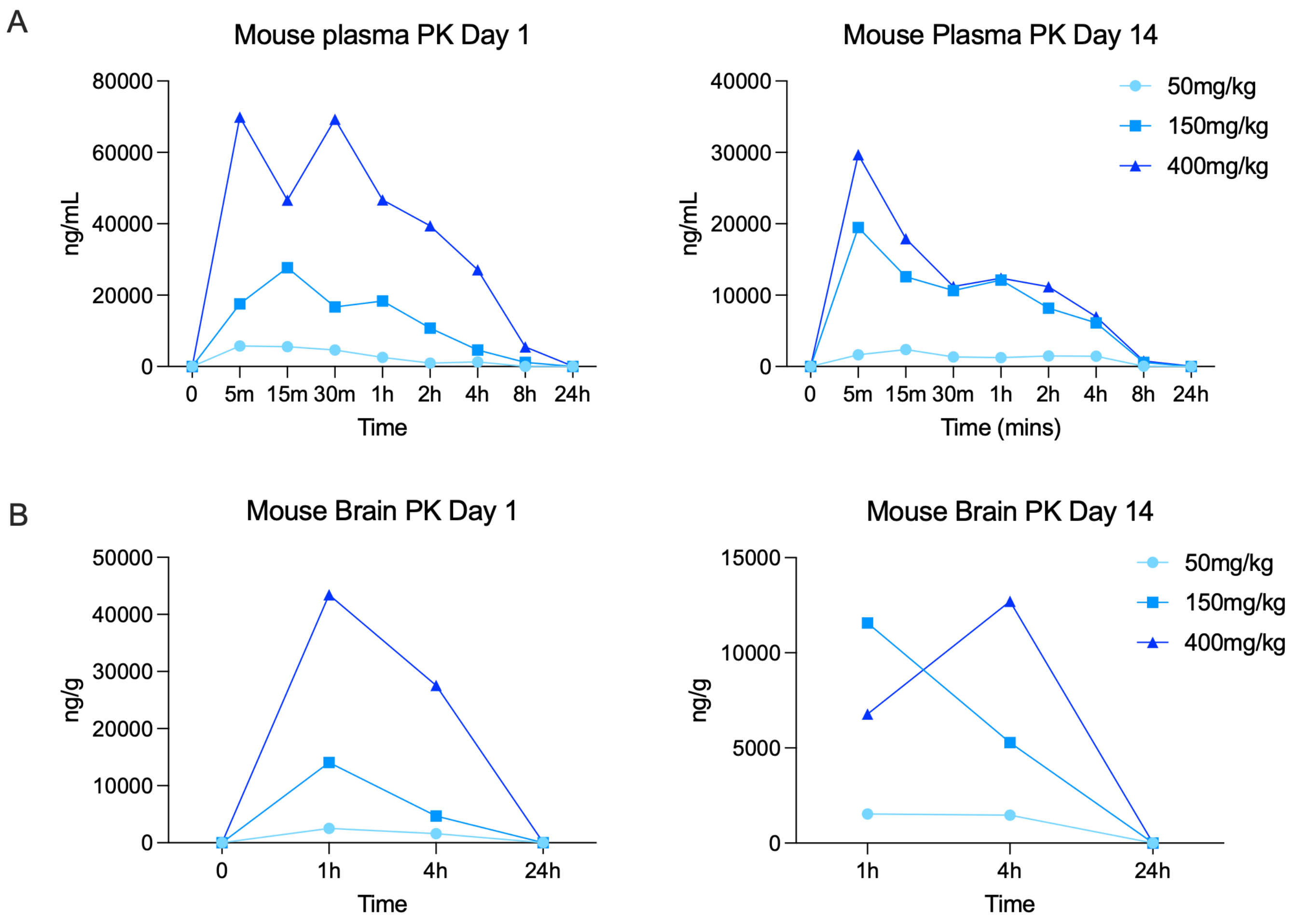

**Figure S7. ACE-339 does not affect TDP-43 expression, microgliosis and additional behavioral phenotypes in iTDP43$^{A315T}$ mouse model.** (A-C) Immunohistochemistry for hTDP-43, microglia and cortical neurons in mouse brain sections. ACE-339 treatment does not affect the elevated hTDP-43 (A) and Iba1 (B) levels in iTDP43 mice cortex and hippocampus. No changes were observed in Ctip$^{+}$ cortical neurons in the model or with ACE-339 treatment. (D) ACE-339 treatment does not affect rotarod motor behavior phenotype. (E-F) ACE-339 treatment shows reduction in time to descend in motor pole test (E) and time spent in open arm in elevated plus maze (F). (G) ACE-339 does not significantly affect distance travelled and time spent in the center of an open field test. Mean±SEM are shown. $P<0.05$*, $P<0.01$**, $P<0.001$***, $P<0.0001$**** One-way ANOVA followed by Kruskal-Wallis test.

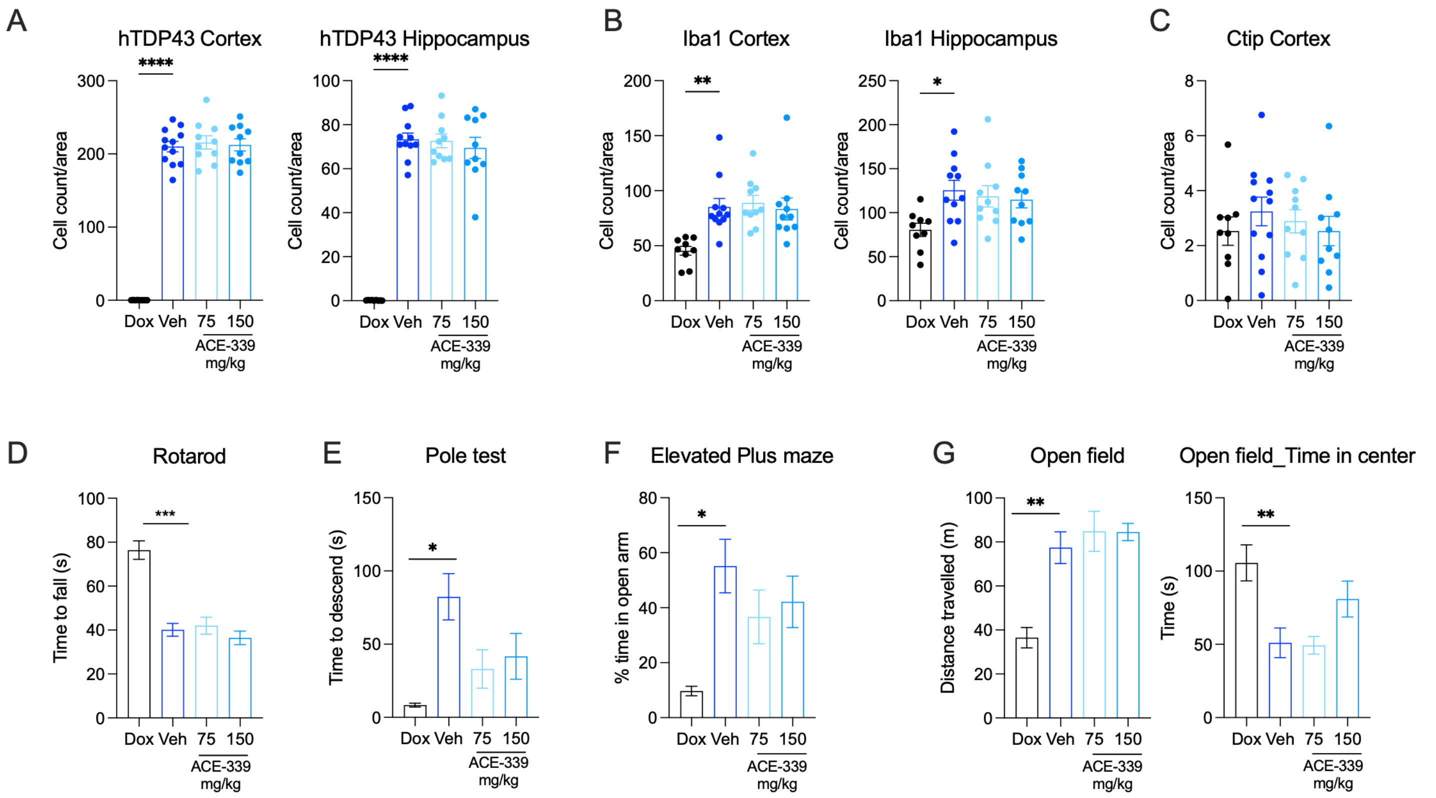

**Figure S8. ACE-339 shows no significant gender differences in rNLS8 mice.** (A) Survival curve separated by gender shows that ACE-339 improved survival in both female and male rNLS8 mice. (B) No significant gender differences observed in plasma Nf-L after 3 and 4 weeks of ACE-339 dosing. (C) Body weights of mice in each treatment group over the entire study period. (D) Plasma and brain ACE-339 drug level measurement (PK) in an independent satellite group of rNLS8 mice (n=4-5/group) treated as described in Figure 3F after 4 weeks of dosing. (E) Biochemical fractionation of insoluble hTDP-43 from rNLS8 mouse brains measured with Simple Western analysis. Following disease induction and subsequent treatment, ACE-339 reduced the accumulation of pathological, insoluble hTDP-43 in a dose-dependent manner, demonstrating a strong trend toward clearance at the highest dose (p = 0.0506) compared to vehicle controls. Mean±SEM are shown in B-C, unpaired t-test in E.

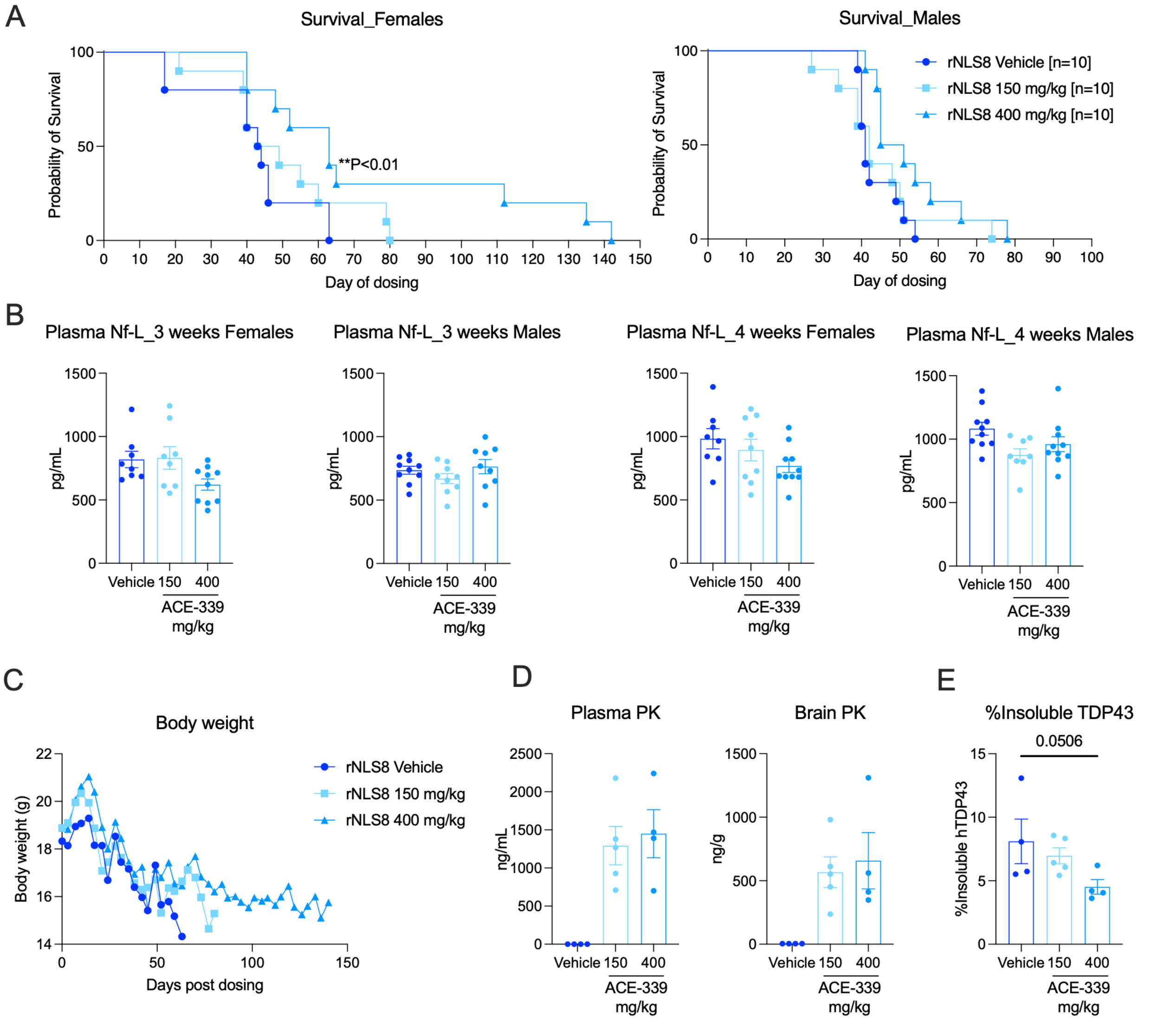

**Figure S9. ACE-339 drives a transcriptome-wide isoform expression shift in disease pathways in rNLS8 mice**. (A) Volcano plot of global differential isoform expression showing shifts in ACE-339 treated rNLS8 brains compared to vehicle controls. Genes containing the top 20 significant differentially expressed isoforms are highlighted. (B) Pathway enrichment analysis of genes with significantly depleted isoforms in response to ACE-339, highlighting impacts on of cell death, stress, and inflammatory responses. (C) Pathway enrichment analysis of genes with significantly increased isoforms in response to ACE-339, demonstrating impacts on neurogenesis and neurodevelopmental networks. Cutoffs: p-adj < 0.05 & |log2FC| >1.

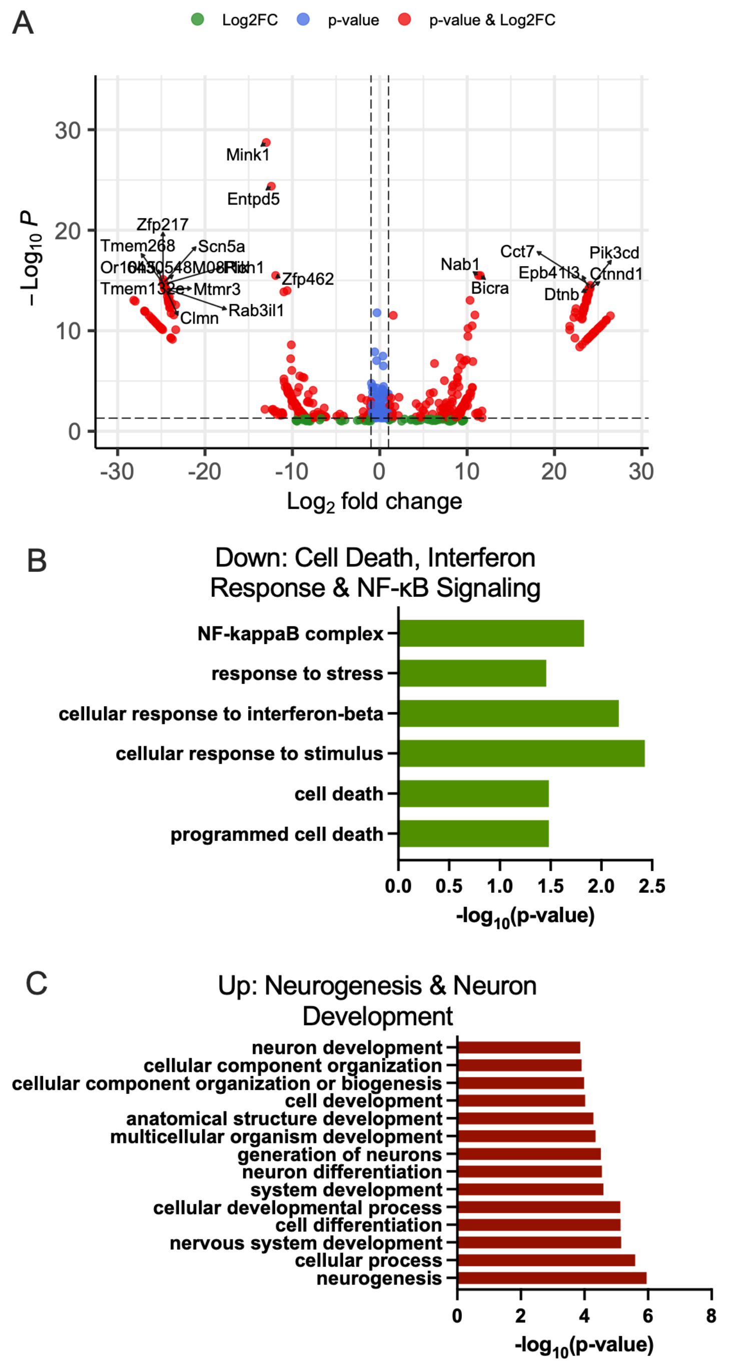

**Figure S10. Total TDP-43 concentration in ALS post-mortem brain, iTDP43$^{A315T}$ brain and rNLS8 brain tissues.** Human tissue was the motor cortex region of the brain. Mouse models were without any drug treatment. Numbers on the top of each bar are the respective means of the group.

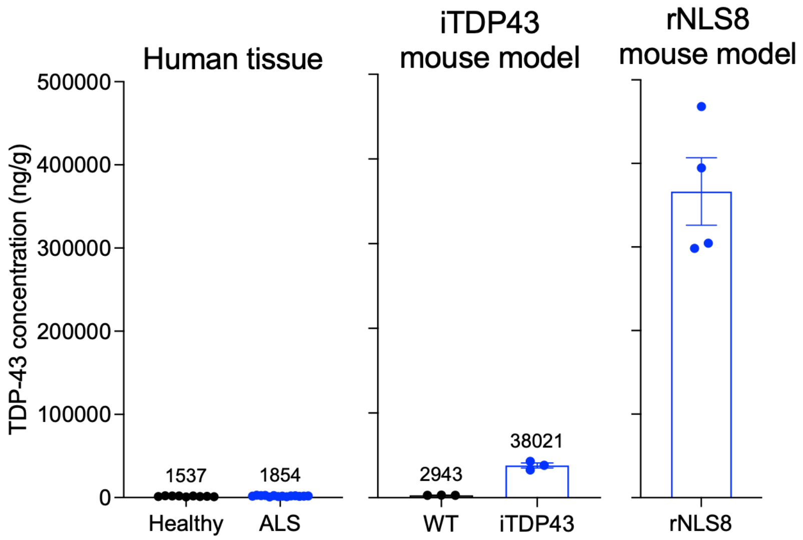

**Figure S11. Molecular dynamics (MD) simulations show that ACE-339 interacts with TDP-43 at specific residues during disaggregation.** (A) The interpeptide contact map from the replica exchange MD simulation of the TDP-43 (307-319) hexamer in in aqueous conditions, demonstrating the close proximity between phenylalanine side-chains. (B) Chemical groups 1-5 within the ACE-339 small molecule interact with the residues on the TDP-43 (307-319) hexamer with the frequency of those peptide-ligand contacts measured. (C) The number of ACE-339 molecules in contact with the TDP-43 (307-319) hexamer over the course of periods within the simulation, averaged over the first 300 ns (left), the following 300 to 600 ns (right).

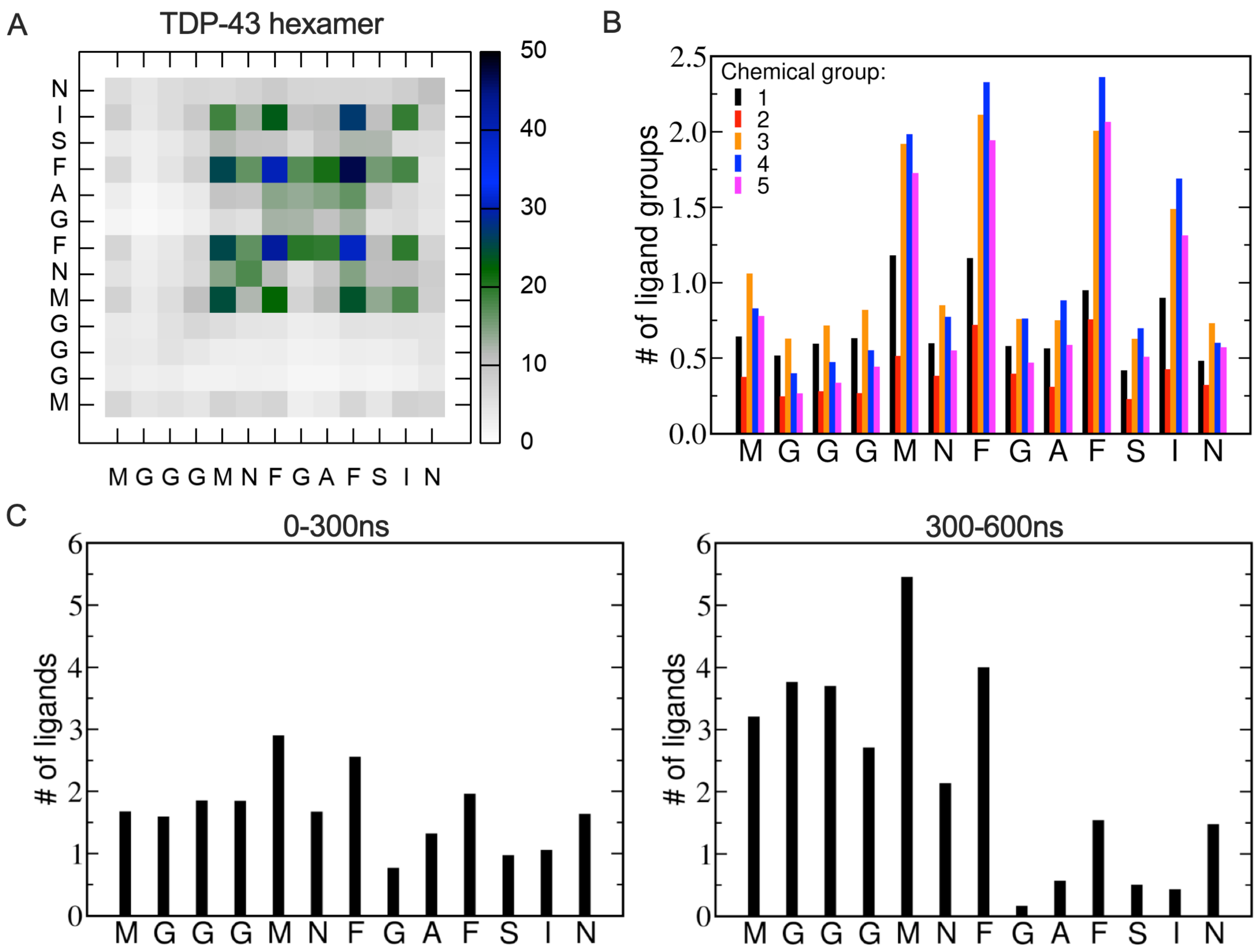

**Table S1. Disease and additional details of human brain donors used in this study.**

| Group | Sample ID | Source | Sex | Age | PMI (h) | TDP-43 pathology status | Additional information |
|---|---|---|---|---|---|---|---|
| Non-neurological control | GBB23-11 | Georgetown University | Male | 62 | 36 | | |
| Non-neurological control | GBB-21-03 | Georgetown University | Male | 71 | 47 | | Diabetes, hypertension, chronic kidney disease, bronchiectasis |
| Non-neurological control | GBB-23-12 | Georgetown University | Male | 67 | 35 | | Hypertension |
| Non-neurological control | GBB-21-27 | Georgetown University | Female | 82 | 27 | | Diffuse large cell lymphoma, hypertension, Fungal mycosis |
| Non-neurological control | SD036/18 | University of Edinburgh | Female | 49 | 69 | Control, No significant abnormalities | |
| Non-neurological control | SD034/15 | University of Edinburgh | Male | 69 | 49 | Control, No significant abnormalities | |
| Non-neurological control | SD035/14 | University of Edinburgh | Female | 59 | 53 | Control, No significant abnormalities | |
| Non-neurological control | SD005/14 | University of Edinburgh | Male | 63 | 42 | Control, No significant abnormalities | |
| sALS | GWF-15-08 | Georgetown University | Female | 61 | 3.5 | sALS, high TDP inclusion frequency | |
| sALS | GWF-15-12 | Georgetown University | Male | 74 | 7 | sALS, early AD, high TDP inclusion frequency | |
| sALS | GWF-15-10 | Georgetown University | Female | 63 | 37.5 | sALS, high TDP inclusion frequency | |
| sALS | GWF-15-14 | Georgetown University | Male | 74 | 3.5 | sALS, high TDP inclusion frequency | |
| sALS | SD027/24 | University of Edinburgh | Male | 64 | 42 | Motor Neuron Disease - ALS-TDP pathology, Thal amyloid phase 1, Braak tangle stage 1, WM pathology: mild, Mild non-amyloid SVD | Limb onset |

| | | | | | | | |
|---|---|---|---|---|---|---|---|
| sALS | SD019/24 | University of Edinburgh | Male | 50 | 24 | Motor Neuron Disease - ALS-TDP pathology, Mild arteriolar Aβ-CAA, Mild non-amyloid SVD | |
| sALS | SD008/24 | University of Edinburgh | Male | 73 | 88 | Motor Neuron Disease - ALS-TDP pathology, Thal amyloid phase 3, Braak tangle stage II, Lewy Body Disease (LBD): brain stem subtype, WM pathology: moderate, Moderate non-amyloid SVD, Moderate/severe arteriosclerosis in occipital white matter | Respiratory onset |
| sALS | SD002/24 | University of Edinburgh | Male | 77 | 70 | Motor Neuron Disease - ALS-TDP pathology, Braak tangle stage II, WM pathology: moderate, Moderate non-amyloid SVD, FTLD-TDP Type A | Dementia onset |
| sALS | SD027/22 | University of Edinburgh | Female | 71 | 91 | Motor Neuron Disease - ALS-TDP pathology, Thal amyloid phase 1, Braak tangle stage 1, WM pathology: severe, Severe non amyloid SVD | Bulbar onset |
| sALS | SD045/21 | University of Edinburgh | Female | 53 | 59 | Motor Neuron Disease - ALS-TDP pathology, WM pathology: mild, Mild non-amyloid SVD | Unknown |
| sALS | SD021/21 | University of Edinburgh | Female | 61 | 45 | Motor Neuron Disease - ALS-TDP pathology, Braak tangle stage 1, WM pathology: mild, Mild non-amyloid SVD | Bulbar onset |
| sALS | SD006/21 | University of Edinburgh | Male | 40 | 68 | Motor Neuron Disease - ALS-TDP pathology, WM pathology: mild, Mild non-amyloid SVD | |

| | | | | | | | |
|---|---|---|---|---|---|---|---|
| sALS | SD018/20 | University of Edinburgh | Female | 59 | 44 | Motor Neuron Disease - ALS-TDP pathology, WM pathology: mild, Mild non-amyloid SVD, FTLD-TDP-43 | Bulbar onset |
| sALS | SD008/20 | University of Edinburgh | Male | 72 | 44 | Motor Neuron Disease - ALS-TDP pathology, Braak tangle stage I, WM pathology: mild, Mild non-amyloid SVD, FTLD-TDP-43 | Limb onset |
| ALS | | Institute for Brain Aging and Dementia at UC Irvine | Male | 64 | 17 | Braak tangle stage I | |
| ALS | | Institute for Brain Aging and Dementia at UC Irvine | Male | 68 | 24 | Braak tangle stage I | |
| ALS | | Institute for Brain Aging and Dementia at UC Irvine | Male | 84 | 30 | Braak tangle stage 0 | |

**Table S2. List of the qRT-PCR primers used in this study.**

| Gene | Forward primer (5'-3') | Reverse primer (5'-3') |
|---|---|---|
| GAPDH | GAAGGTGAAGGTCGGAGTC | GAAGATGGTGATGGGATTTC |
| TDP43 | GATGGTGTGACTGCAAACTTC | CAGCTCATCCTCAGTCATGTC |
| STMN2 | AGCTGTCCATGCTGTCACTG | GGTGGCTTCAAGATCAGCTC |
| KALRN | TTAGTCCCGGGTGTCCTTAT | AGCAGCATCATATTCTGGAAGT |
| UNC13A | GGACGTGTGGTACAACCTGG | GTGTACTGGACATGGTACGGG |
| STMN2 cryptic exon | GCTCTCTCCGCTGCTGTAG | CTGTCTCTCTCTCTCGCACA |
| KALRN cryptic exon | TCCAGAATATGGCTGCTCACT | CAGCAGCATCTTGTGTGATGT |
| UNC13A cryptic exon | TGGATGGAGAGATGGAACCT | GGGCTGTCTCATCGTAGTAAAC |
| CDK7 cryptic exon | GCAGTGTGGACATGACTGATA | GTCCACACCTACACCATACATC |
| ACTL6B cryptic exon | ATCCTGGATCACACCTACAGC | AGGAGGATTGCTTGAACCC |
| CAMK2B cryptic exon | GAGTGCAGAGACTTCCCCC | CTGCTCCGTGGTCTTAATGAT |
| HDGFL2 cryptic exon | TCACACCTGAGAAGAAAGCAG | TCCTCTCTTCTGTGTCCCTCT |
| MYO18 cryptic exon | AAGTCCAGGGATGAGATTGTG | GCAGAGTTTTGTCCTCCTCTTTA |
| SYT7 cryptic exon | GCAGTGAGAAGAAGGCTATCAA | CGGCAGACTGGAGCCT |
